# Environmental drivers of the resistome across the Baltic Sea

**DOI:** 10.1101/2025.03.06.641941

**Authors:** Joeselle M. Serrana, Francisco J. A. Nascimento, Benoît Dessirier, Elias Broman, Malte Posselt

## Abstract

**Background:** Antimicrobial resistance is a major global health concern, with the environment playing a key role in its emergence and spread. Understanding the relationships between environmental factors, microbial communities, and resistance mechanisms is vital for elucidating environmental resistome dynamics. In this study, we characterized the environmental resistome of the Baltic Sea and evaluated how environmental gradients and spatial variability, alongside its microbial communities and associated functional genes, influence resistome diversity and composition across geographic regions.

**Results:** We analyzed the metagenomes of benthic sediments from 59 monitoring stations across a 1,150 km distance of the Baltic Sea, revealing an environmental resistome comprised of predicted antimicrobial resistance genes (ARGs) associated with resistance against 26 antibiotic classes. We observed spatial variation in its resistance profile, with higher resistome diversity in the northern regions and a decline in the dead zones and the southern areas. The combined effects of salinity and temperature gradients, alongside nutrient availability, created a complex environmental landscape that shaped the diversity and distribution of the predicted ARGs. Salinity predominantly influenced microbial communities and predicted ARG composition, leading to clear distinctions between high-saline regions and those with lower to mid-level salinity. Furthermore, our analysis suggests that microbial community composition and mobile genetic elements might be crucial in shaping ARG diversity and composition.

**Conclusions:** We presented that salinity and temperature were identified as the primary environmental factors influencing resistome diversity and distribution across geographic regions, with nutrient availability further shaping these patterns in the Baltic Sea. Our study also highlighted the interplay between microbial communities, resistance, and associated functional genes in the benthic ecosystem, underscoring the potential role of microbial and mobile genetic element composition in ARG distribution. Understanding how environmental factors and microbial communities modulate environmental resistomes will help predict the impact of future environmental changes on resistance mechanisms in complex aquatic ecosystems.

## Background

Antimicrobial resistance is a global health concern that involves the transmission of pathogens and genes across animals, humans, and environmental systems [1]. Recently, the role of the environment in the emergence and transmission of antimicrobial resistance has been increasingly recognized [2–4]. Microorganisms acquire resistance through de novo mutation or by horizontal gene transfer of antimicrobial resistance genes (ARGs) encoded on mobile genetic elements (MGEs, e.g., plasmids, integrons, or transposons) between similar or different species [5]. Resistant microorganisms and ARGs are then transmitted across environments and species boundaries, demonstrating the need to understand the connections between antimicrobial resistance and environmental microbiomes [6, 7].

The environmental resistome – the collection of all types of acquired and intrinsic resistance genes [8–10] in aquatic ecosystems is receiving growing interest. This also includes precursor resistance genes and latent resistance mechanisms within microbial communities that may require evolutionary changes or shifts in expression context to confer resistance [8]. Rivers, lakes, and marine coastal habitats have higher resistance burdens, where pathogens and ARGs spread and are potentially selected due to exposure to environmental contaminants [11–13]. Aquatic ecosystems receive inputs from various anthropogenic sources, e.g., pharmaceutical and other polluting industries, increasing their resistance loads through various mechanisms (e.g., increasing horizontal gene transfer rates) [4]. Furthermore, environmental factors also influence the emergence and dissemination of resistance in natural environments [14]. Several reports measured and assessed the direct and indirect effects of environmental factors as stressors on environmental resistomes to assess their relationship and explain underlying resistance mechanisms [15–19]. However, no consensus exists on which environmental factor mainly drives antimicrobial resistance in complex aquatic ecosystems [15, 20]. Mainly because multiple environmental factors would have cumulative or synergistic effects that influence variable dynamics in the resistomes of different environments. Given that multiple environmental factors contribute to the maintenance and proliferation of resistant microorganisms and ARGs in natural environments, an improved understanding of the environmental drivers of the resistome in potential natural reservoirs would be relevant for antimicrobial resistance surveillance and monitoring, necessary for predicting resistance dissemination and the development of effective management strategies.

Profiling the environmental resistome and its mobilization are also critical to understanding the role of the environment as a major source of resistance factors [2, 9]. However, the internal link between ARGs and their co-selection with metal resistance genes (MRGs), microbial community composition, and the functional genes involved in its mobility and host pathogenicity, i.e., MGEs and virulence factor gene (VFGs), remain underexplored. In particular, there is limited understanding of the connection between resistance and VFGs, which play a crucial role in the infectivity of pathogenic organisms. Direct and co-selection may occur over a wide concentration range of pollutants and physicochemical gradients [4, 21]. We still lack comprehensive information on the pathways of antimicrobial agents in the environment, as well as their accumulation, persistence, and effective concentration. Hence, characterizing the resistome and understanding the link between ARGs, resistance-associated functional genes, and their environmental transmission remains complex and challenging due to these multiple knowledge gaps.

The Baltic Sea is a large brackish coastal ecosystem with pronounced environmental gradients that primarily govern its biodiversity and benthic communities [22–33]. These gradients are created due to the sea’s semi-enclosed location, alongside strong influences from the surrounding landmasses and climate [25]. Due to the strong environmental gradients in a relatively short spatial scale, e.g., salinity, depth, nutrient loads, temperature, and O_2_ conditions [22, 26], the Baltic Sea is an ideal ecosystem for assessing the interaction between environmental resistomes and their microbial communities to fully understand the mechanisms of resistance in complex environments. In particular, the Baltic Sea’s North-to-South gradients from freshwater to brackish and marine conditions make it an interesting ecosystem for studying resistomes due to its distinct ecological niches and differential microbial diversity [24–27]. These large gradients have the potential to enhance our understanding of how varying environments influence the development and spread of resistant microbial communities in response to environmental change.

In this study, we analyzed the metagenomics data from benthic sediments collected from 59 monitoring stations spanning 1,150 km distance of the Baltic Sea [27]. We (i) characterized the environmental resistome, i.e., ARGs and MRGs, of the benthic ecosystem alongside its microbial communities and specific functional genes, i.e., MGEs and VFGs, (ii) investigated their correlations, and (iii) assessed how environmental gradients and spatial variability influence the diversity and composition of the resistome across geographic regions. We hypothesize that similar to the observed distribution of benthic functional genes and metabolic pathways by Broman et al. [27], there were also regional differences in the resistome and its associated functional genes and that ARG diversity and composition were mainly structured by specific environmental factors. Additionally, given the complex relationships between microbial communities and their resistance mechanisms, we hypothesize that microbial community composition directly and indirectly influenced the resistome. To our knowledge, this is the first report characterizing the environmental resistome of the Baltic Sea and its association with the benthic sediment microbiome’s mobility and virulence capacities. Lastly, we provided key insights regarding the environmental resistome of a model ecosystem with distinct physicochemical gradients and potential selective factors, highlighting the critical role of the environment in the selection and spread of antimicrobial resistance in complex aquatic ecosystems.

## Methods

### Sample description

Broman et al. [27] generated the metagenomics data of benthic sediment samples collected by the Swedish National and Regional Benthic Monitoring Program in the Baltic Sea. Between May and June 2019, 59 benthic sediment samples were collected from selected soft-bottom clay-muddy habitats from the north to the southern regions of the Baltic Sea (**Supplementary Table S1**). The sediments were collected via core sampling, and the top 2 cm layer of the samples was sliced and preserved at −20°C in the field. After homogenization, 0.2 g of this preserved sediment was used for total genomic DNA extraction, followed by metagenomics sequencing as described in Broman et al. [35]. Environmental parameters measured in this study include water depth (m), salinity, temperature, and dissolved O_2_ (mg/L). The concentrations of total carbon (TC) and total nitrogen (TN) (µmol/g), and their ratio – C/N, and isotopic signatures, i.e., stable δ^13^C and δ^15^N isotope compositions (‰) were also assessed as indicators of resource availability which has crucial impacts on microbial community dynamics by influencing competition, metabolic pathways, and responses to environmental changes. In particular, the stable isotope composition provides an idea of the origin of organic matter deposited in the sediments (**Supplementary Table S1** and **Fig. S1**). The samples were grouped into eight regions (**Fig. 1A**), i.e., Bothnian Bay, Bothnian Sea, Stockholm, Sörmland, Dead Zone (Sörmland), Östergötland, Dead Zone (Mid-South), and Southern Baltic, based on the geographic locations and basin bathymetry of the monitoring stations from which they were collected [26–27].

**Fig. 1.**
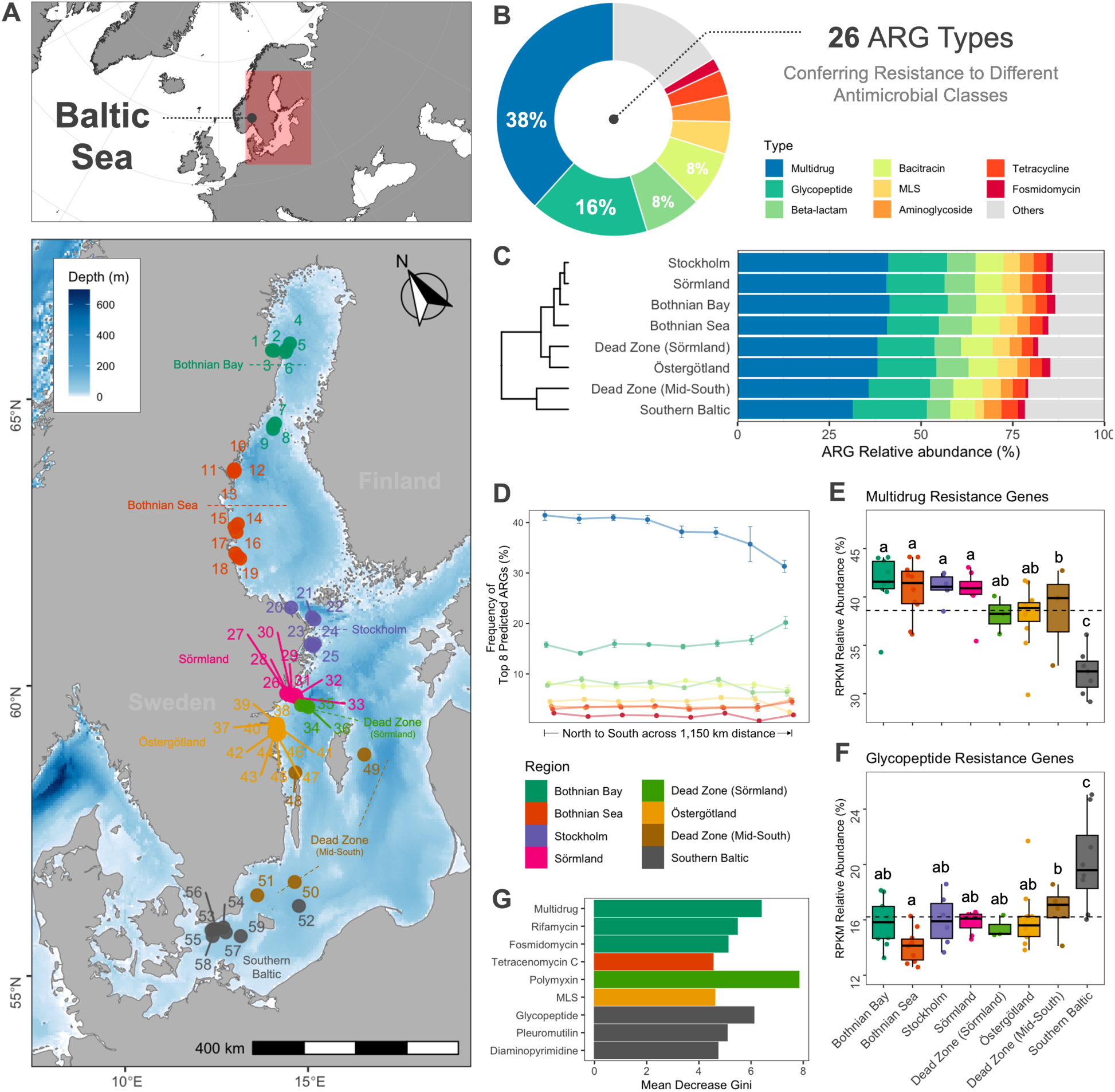
Resistome composition and diversity across the Baltic Sea. **A** The map of the sampling stations along 1,150 km across the Baltic Sea grouped by spatial location as described in Broman et al. (2022). **B** Relative abundance of the top 8 ARG types identified in the study. **C** Average relative abundance of the ARG types per region and hierarchical clustering. **D** Relative frequency of the top 8 ARG types per region. Line colors correspond to the ARG types in **Fig. 1B**. Relative abundance of the **E** multidrug and **F** glycopeptide resistance types by region. Different letters indicate differences between groups tested by ANOVA (*P*-value < 0.05). See Fig. S4A for the relative abundances of the top 10 ARG types. The dotted horizontal line indicates the mean RPKM relative abundance. **G** Differential abundance test by Kruskal-Wallis rank sum and random forest. The variable importance plot shows the differentially abundant ARG types (adjusted *P*-value < 0.05). See **Fig. S5B** for the variable importance plot of the top ARG subtypes with the highest mean decrease Gini values.

### Metagenomics data processing

#### Read processing

Raw sequencing data with an accession number PRJEB41834 was recovered from the European Nucleotide Archive (ENA) database. Sample identifiers and metadata were obtained from the supplementary tables of the published paper [27]. The raw sequence reads were processed using fastp v0.23.4 [28] to remove adaptor sequences and quality filter paired-end reads using default parameters (i.e., phred quality >= Q15). Human and other contaminant sequences, i.e., PhiX, were removed by mapping the reads to a non-redundant version of the Genome Reference Consortium Human Build 38 (GRCh38hg38; RefSeq: GCF_000001405.40) and the phiX reference genomes (GCF_000819615.1) performed with the Kraken2 v2.1.2 package [29]. The quality-filtered paired-end reads were error-corrected using bbcms v38.61b from the BBTools [30; https://sourceforge.net/projects/bbmap/] with default parameters.

#### Contig assembly, gene prediction, and read mapping

Individual assemblies were performed with MEGAHIT v1.2.9 [31] using the preset parameters for the meta-large option (--k-min 27 --k-max 127 --k-step) as recommended in the MEGAHIT documentation for complex metagenomics data. Genes were predicted on the assembled contigs using the PROkaryotic Dynamic programming Gene-finding ALgorithm (Prodigal) v2.6.3 [32]. The predicted open reading frames (ORFs) or protein-coding genes were translated into amino acid sequences using the anonymous gene prediction mode (prodigal -p meta) and default parameters. Read counts of each gene were estimated by mapping the quality-filtered paired-end reads to the contigs with minimap2 v2.24-r1175 [33; https://github.com/lh3/minimap2] and summarizing the read abundance of the genes into count tables using featureCounts v2.0.3 [34]. See **Supplementary Table S2** for a detailed count list for each metagenomic data processing per sample.

### Taxonomic profiling of microbial species

A reference genome-independent species-level profiling of metagenomes based on single-copy phylogenetic marker gene sequences was performed with the metagenomic Operational Taxonomic Units (mOTU) profiler v3.0.2 [35; see https://github.com/motu-tool/mOTUs]. The tool estimates the relative taxonomic abundance of known and currently unknown microbial community members from metagenomic sequencing data. With the quality-filtered paired-end reads as input, taxonomic profiling was performed with the *motus profile* command set to default parameters. Each sample’s identified species and relative abundance information were merged into one mOTU table for the downstream analysis (**Supplementary Table S3**).

### Annotation of resistance genes, mobile genetics elements, and virulence factors

#### Prediction of antimicrobial resistance genes

Following contigs assembly and prediction of protein-coding regions, ARGs were annotated using different computational approaches and against different ARG databases (**Supplementary Table S4**). DeepARG v.2.0 [36] was chosen as the ARG annotation method due to its higher detection rate and better performance in the environmental association analyses, identifying 2,042 annotations (belonging to 378 ARGs) with a dbRDA adjusted *R^2^* value of 0.24, compared to the other methods or databases, which detected fewer annotations despite having similar or lower *R²* values (**Fig. S2**). These indicate that the ARG composition based on DeepARG demonstrated stronger environmental association compared to other methods, providing a better understanding of regional variations, distance decay relationships, and the impact of specific environmental factors on the resistome in the Baltic Sea.

DeepARG uses a deep learning-based approach to annotate ARGs, which improves annotation accuracy, especially for genes with low sequence similarity to the ARG reference. In this study, DeepARG annotation was performed from the predicted protein sequences (translated Open Reading Frame sequences; ORFs) with the DeepARG-LS model against the DeepARG database. The model provides more accurate antimicrobial resistance annotations compared to the traditional best-hit approach, resulting in lower false-negative rates and higher overall reproducibility [37]. It consists of four dense hidden layers that propagate a bit score distribution, and the output layer of the deep neural network corresponds to 102 antibiotics from 30 antimicrobial resistance categories. AMR categories refer to the functional potential of the ORFs associated with the encoded resistance. The DeepARG model was trained against three curated databases, i.e., the Comprehensive Antibiotic Resistance Database (CARD) [38], Antibiotic Resistance Genes Database (ARDB) [39], and UniProt [40]. Default options were used (deeparg --model LS --min-prob 0.8 --arg-alignment-evalue 1e-10) as Arango-Argoty et al. [36] recommended. We note that DeepARG predicts ARG classes, which include genes that are not inherently ARGs but are associated with antimicrobial resistance (e.g., transporter genes), which are still crucial in a resistome analysis. Predicted ARG abundances were expressed as reads per kilobase million (RPKM) (**Supplementary Table S5**). RPKM was used to normalize gene coverage values by correcting the differences in sample sequencing depth and gene length. The RPKM value of the gene is calculated using the equation [41]: *RPKM* = *numReads*/((*geneLength*/10^3) × (*totalNumReads*/10^6)), where *numReads* is the number of reads mapped to a gene sequence, *geneLength* is the length of the gene sequence, and *totalNumReads* is the total number of mapped reads of a sample. The relative ARG abundance across samples was visualized by summing the RPKM for all ARGs belonging to a type (AMR category) or a subtype (gene level).

#### Screening potential metal resistance genes

The protein-coding genes were also matched against the BacMet v.2.0 predicted resistance genes database [42; http://bacmet.biomedicine.gu.se/] with DIAMOND v2.1.6 [43] to screen for potential MRGs from the samples. The BacMet database consists of high-quality and manually curated bacterial genes that have been experimentally confirmed to confer resistance to metals and antibacterial biocides with full references in the scientific literature [42]. DIAMOND was performed with the blastp command using the following flags: --sensitive -e 1e-10 -k 1 --id 80 --query-cover 70. The abundance of each MRG was expressed in RPKM (**Supplementary Table S6**). The relative MRG abundance across samples was visualized by summing the RPKM at the gene level.

#### Annotation of mobile genetic elements and virulence factors

Identification and annotation of MGEs and VFGs were performed by searching the protein-coding gene sequences against the mobileOG-db release beatrix-1.6 [44; https://github.com/clb21565/mobileOG-db], and the virulence factor database (VFDB) set B [45; http://www.mgc.ac.cn/VFs/download.htm] with the diamond blastp command using similar probability values set for the MRG screening. The mobileOG-db (mobile orthologous groups database) is a manually curated database of protein families that mediates integration/excision, replication/recombination/repair, stability/defense, or transfer of bacterial mobile genetic elements and phages, as well as the associated transcriptional regulators of these processes [44]. VFDB is a collection of bacterial virulence factor protein families with in-depth coverage of the major virulence factors of relevant pathogenic bacteria [45]. It contains functionally classified fasta files of bacterial virulence factor protein sub-families. The abundance of each identified MGE and VFG across samples was also expressed in RPKM (**Supplementary Table S7** and **S8**). The relative abundances of MGEs and VFGs across samples were visualized by summing the RPKM at their category level.

### Statistical analyses

The statistical analyses and visualizations were performed in the R environment v4.4.2 [46]. The Baltic Sea and sample location map was produced using the ggOceanMaps v2.2.0 package [47]. Autocorrelations among the environmental factors were visualized in correlograms (**Supplementary Fig. S3**) and tested for multicollinearity by calculating the variance inflation factor (VIF) using the vif function in R. A VIF < 5 was used as the selection criteria, hence, the concentrations of TN and TC (%) were excluded in the downstream analyses as they were collinear with the isotopic signatures, i.e., δ13C and δ15N. The geographic distance for each site (in km) was measured using the haversine formula, calculated from the latitude and longitude coordinates of each site relative to the northernmost site (i.e., S01) using the geosphere package [48]. The microeco v1.13.0 [49] package was used to calculate the alpha and beta diversities of the microbial community and the gene datasets, i.e., ARGs, MRGs, MGEs, and VFGs. The alpha diversity indices assessed were Chao1 richness, Shannon–Wiener diversity, and Pielou’s evenness. The abundance-based Bray-Curtis dissimilarity index was used to assess beta diversity, which was estimated using relative abundances for the microbial community data and RPKM values for the gene datasets. Principal coordinate analysis (PCoA) was performed to visualize the Bray-Curtis dissimilarity matrices. To assess the significant differences in alpha and beta diversity metrics between regions, a one-way analysis of variance (ANOVA) with Duncan’s multiple comparison tests or the Dunn’s Kruskal-Wallis (KW) multiple comparisons test was used. For this, the homogeneity of variances was checked using Levene’s test with the car package [50]. A non-parametric multivariate analysis of variance (PERMANOVA) was used to infer significant differences in community composition based on Bray-Curtis distance among the sampled regions. Both were assessed at a significant level of *P-value* < 0.05. To assess the homogeneity of multivariate dispersions of the Bray-Curtis dissimilarity matrices, we used the betadisper function from the vegan v2.6.6 package in R [51]. Linear regression was performed to assess the distribution of the alpha diversity estimates of the 59 samples along the 1,150 km distance of the Baltic Sea using the general linear model function in R. The distance decay rate of ARGs was plotted using the plot_scatterfit function of the microeco package. Distance decay was computed as the slope of the ordinary least-squares regression line fitted to the relationship between the haversine (km) distance and the Bray-Curtis dissimilarity distance.

A random forest analysis coupled with a differential abundance test [52] was carried out on the predicted ARG dataset based on RPKM using the trans_diff class of the microeco package to test differentially abundant ARG types and subtypes among the sampled regions. The algorithm was performed with regions as response variables, and the root variables for building a random forest of classification trees were set by default with 1,000 grown trees. A similar approach was performed on the mOTU dataset to identify differentially abundant microbial species among the regions. The random forest approach was limited to detecting significantly abundant predicted ARGs, without accounting for the complex interactions and dependencies among different types and subtypes, as the method does not explicitly model these interactions. A partial least squares path modeling (PLS-PM) analysis was performed using the plspm v0.5.1 package [53] to distinguish the direct and indirect relationships between the microbial communities and their gene profiles. PLS-PM is a correlation-based structural equation modeling algorithm that estimates complex cause-effects or prediction models using latent variables. Before the analysis, the microbial community and gene datasets were filtered by prevalence set to 0.05. Bootstrap validation was set to 1,000, with the other parameters set to default. For the inner model path matrix, the microbial community was set as the exogenous variable acting as the primary predictor, influencing MGEs, which in turn, along with microbial communities, influence VFGs, ARGs, and MRGs; however, MRGs are not directly influenced by ARGs. Results of the inner model were plotted, including the path coefficients and *R^2^* values for each latent variable. The PLS-PM analysis does not capture complex interactions or non-linear relationships, limiting its ability to fully reflect intricate dependencies in the data. Hence, a significance test for the path coefficient and loading factor was performed using a t-test, and Mantel tests and Procrustes analysis were used to evaluate the congruency between samples based on their microbial communities and resistance gene profiles using Bray-Curtis dissimilarity matrices with the vegan package. Prior to the test, the dimensionality of the datasets was reduced through principal coordinate analysis.

Pearson’s correlation analysis was used to identify the initial linear relationships between the environmental factors and the alpha diversity indices. To further explore non-linear relationships and assess multiple predictors and interactions, generalized additive models (GAMs) were used to examine the effects of environmental factors on predicted ARG richness, diversity, and evenness. The GAMs were constructed using the mgcv package [54] with the alpha diversity metrics (i.e., Chao1, Shannon, and Pielou) as response variables and each environmental factor as predictor through smooth functions. The association between environmental factors and predicted ARG composition was assessed using distance-based redundancy analysis (dbRDA) using the trans_env class in the microeco package. A random forest analysis was also performed to identify the relevant contributions of the environmental factors on predicted ARG composition. The Euclidean distances between samples for each environmental factor were estimated using the trans_env class, and scatter plots were used to visualize and calculate Pearson’s correlation between the predicted ARG profile (Bray-Curtis distance) and the Euclidean distance of each environmental factor. Spearman’s correlation analysis was also performed to assess the relationships between the environmental factors and ARG types using the microViz v0.12.1 package [55].

## Results

### Environmental resistome profile of the Baltic Sea benthic ecosystem

The predicted antimicrobial resistance profile of the benthic sediments comprised of 26 ARG types (i.e., antimicrobial class) and 378 subtypes (i.e., ARGs and resistance-associated genes). Multidrug resistance was the most dominant ARG type (total RPKM relative abundance of 38.3%), followed by glycopeptide (16.4%), beta-lactam (7.9%), bacitracin (7.6%), macrolide-lincosamide-streptogramin (MLS; 4.5%), aminoglycoside (3.7%), and tetracycline resistance (3.5%) (**Fig. 1B** and **Supplementary Fig. S4A**). The total RPKM relative abundance of the remaining ARG types ranged from 0.02 to 1.8%. Based on the average relative abundance per region, the Bothnian Bay, Bothnian Sea, Stockholm, and Sörmland were clustered together, indicating similar ARG profiles (**Fig. 1C**). The relative frequency of each gene type also showed a similar trend from the northern to the southern regions (**Fig. 1D** and **Supplementary Fig. S5A**).

The northern regions had a higher gene abundance of multidrug resistance (**Fig. 1 E**; ANOVA, *df* = 7, *F* = 8.8, *P* < 0.0001). The dead zones (i.e., Sörmland and Mid-South), Östergötland, and the Southern Baltic displayed unique profiles having notable variations in ARG types and predicted abundances, which could be associated with their distinct environmental condition. These southern regions mainly showed decreased abundance of multidrug resistance genes and increased glycopeptide resistance (**Fig. 1F**; ANOVA, *df* = 7, *F* = 6.6, *P* < 0.0001). Using a random forest analysis coupled with a differential abundance test, we confirmed that the genes conferring multidrug resistance in the Bothnian Bay (Mean Decrease Gini, MDG = 6.4, *P* < 0.01) and glycopeptide resistance in the Southern Baltic (MDG = 6.1, *P* < 0.01) were differentially abundant in these regions (**Fig. 1G** and **Supplementary Table S9**). Additionally, predicted genes conferring resistance to rifamycin (MDG = 5.5, *P* < 0.01) and fosmidomycin (MDG = 5.1, *P* < 0.05) were differentially abundant in the Bothnian Bay. Predicted ARGs conferring resistance to pleuromutilin (MDG = 5.1, *P* < 0.05) and diaminopyrimidine (MDG = 4.7, *P* < 0.01) were differentially abundant in the Southern Baltic. Other ARG types, i.e., tetracenomycin C (MDG = 4.5, *P* < 0.05), polymyxin (MDG = 7.8, *P* < 0.001), and MLS (MDG = 4.6, adjusted *P* < 0.05) resistance, were differentially abundant in the Bothnian Sea, Dead Zone (Sörmland), and Östergötland regions, respectively.

At the subtype level, the most dominant predicted ARG or resistance-associated gene was the multidrug ABC transporter gene (multidrug resistance) - a large gene family with at least 3 million variants, followed by *vanR* (glycopeptide), *bacA* (bacitracin), and *penA* (beta-lactam), with total predicted ARG relative abundances ranging from 5% to 9%. The differential analysis performed to test the significant differences in the distribution of each ARG subtype across the regions revealed that 33 predicted ARGs were significantly abundant at Bothnian Bay, 13 at Bothnian Sea, five at Stockholm, six at Sörmland, 13 at the Dead Zone (Sörmland), six at Östergötland, three at the Dead Zone (Mid-South), and nine at the Southern Baltic (**Supplementary Table S9**). These and the other top 15 most differentially abundant predicted ARGs representing the regions are presented in **Supplementary Fig. S5B**. A notable observation is the presence of the *arnA* gene involved in resistance to polymyxins. This ARG had the highest mean decrease Gini score and was significantly more abundant in the Dead Zone (Sörmland). Also, the multidrug resistance genes *acrB* and *mexH* were differentially abundant in Bothnian Bay, suggesting a notable presence of broad-spectrum resistance mechanisms in the northernmost region.

The Chao1 richness and Shannon diversity of predicted ARGs decreased from the north to the south of the Baltic Sea (**Supplementary Table S10**). Both indices showed a significant decreasing trend along the 1,150 km distance (**Fig. 2A-B**; Chao1 richness, *df* = 58, *F* = 27.3, *P* < 0.0001; Shannon diversity, *df* = 58, *F* = 18.7, *P* < 0.0001). Distance explained 31% and 23% variation (adjusted *R^2^* values) in these two indices. Also, Pielou’s evenness significantly increased along the 1,150 km distance (**Fig. 2C**; *df* = 58, *F* = 4.3, *P* < 0.05). However, only 5% of the variability in evenness was explained by distance. We observed significant regional differences in alpha diversity based on regional groupings, as shown by the multiple comparison test results for Chao1 richness (KW, *df* = 7, *H* = 25.8, *P* < 0.001) and Shannon diversity (KW, *df* = 7, *H* = 23.7, *P* < 0.01) (**Fig. 2D-E** and **Supplementary Table S11**). The Bothnian Bay and Sea regions showed high predicted ARG richness and diversity, with observed counts ranging from 247 to 669 and 346 to 529, respectively. Both regions also exhibited high evenness (**Fig. 2F**: Pielou > 0.93; ANOVA, *Df* = 7, *F* = 3.3, *P* < 0.01), suggesting a balanced predicted ARG distribution. Similarly, Stockholm and Sörmland showed high predicted ARG diversity and evenness, with Shannon diversity ranging from 5.53 to 6.30 and 5.44 to 6.11, respectively, and evenness mostly greater than 0.95. The two dead zones had relatively low diversity in some samples. However, they maintained high evenness, indicating a relatively balanced predicted ARG distribution despite potential environmental stressors in these regions. Östergötland exhibited moderate diversity, with observed predicted ARG counts ranging from 212 to 511 and Shannon diversity between 5.14 and 5.88, with some variations in evenness across samples. Only the Southern Baltic showed significantly lower values than other regions, with Shannon diversity ranging from 3.71 to 5.51. The alpha diversity results indicate that the northern regions, i.e., Bothnian Bay, Bothnian Sea, Stockholm, and Sörmland, have high predicted ARG diversity and abundances. In contrast, the dead zones and the Southern Baltic showed lower predicted ARG diversity, indicating less complex ARG distribution in these regions.

**Fig. 2.**
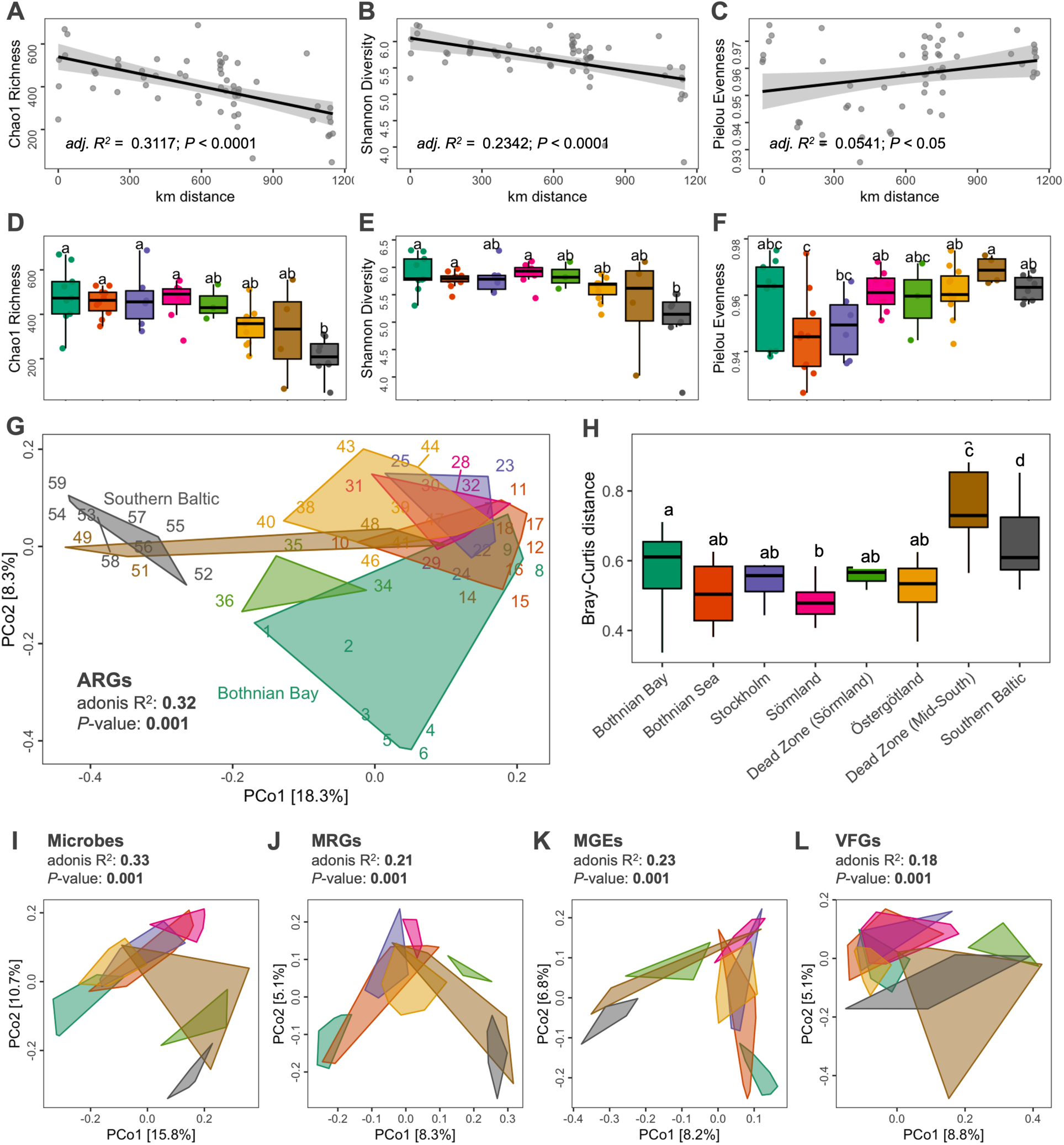
Alpha diversity and composition of resistance and other associated genes. The distribution pattern of predicted ARGs **A** Chao1 richness, **B** Shannon diversity, and **C** Pielou evenness across the Baltic Sea regions in haversine distance (km). **D** Chao1 richness, **E** Shannon diversity, and **F** Pielou evenness per region. Different letters indicate differences between groups tested by ANOVA or Kruskal-Wallis test (*P*-value < 0.05). **G** Principal coordinate analysis (PCoA) plot of the predicted ARG profile based on Bray-Curtis distance grouped by region. **H** Boxplot of the Bray-Curtis distance in each region. Different letters indicate differences between groups tested by ANOVA (p-value <0.05). PCoA plots of the **I** microbial communities, **J** MRG, **K** MGE, and **L** VFG profiles based on Bray-Curtis distance grouped by region.

The principal coordinate analysis (PCoA) based on Bray-Curtis dissimilarity of the predicted ARG profile showed that antimicrobial resistance composition differed between regions, which was statistically confirmed by PERMANOVA (adonis *R^2^* = 0.32; *P* < 0.01; **Supplementary Table S12** and **Fig. 2G**). In particular, the Southern Baltic and some samples from the Dead Zone (Mid-South) were separated from the rest of the sample along the ordination axes. A comparison of within-group distances showed that these two regions have highly dissimilar predicted ARG composition between samples compared to other regions (**Fig. 2H**).

### Benthic microbial composition and functional genetic elements

Using the mOTUs profiler, we identified 4,584 mOTUs, most of which were unique to each sample. A total of 2,772 species belonging to 319 families and 47 phyla were taxonomically annotated. The most abundant phylum was Proteobacteria (relative abundance 51%), followed by Actinobacteria (15%), Bacteroidetes (5%), Thaumarchaeota (4%), and Planctomycetes (4%) (**Supplementary Fig. S4B**). Using a differential analysis, 47 species were significantly differentially abundant among the regions (adjusted *P* < 0.05) (**Supplementary Table S13** and **Fig. S6**). Species with the highest variable importance were *Cyanobium gracile* (Cyanobacteria; Mean Decrease Gini, MDG = 3.3, *P* < 0.001) differentially abundant at the Southern Baltic, *Candidatus Aminicenantes bacterium RBG-13-64-14* (Candidate phylum OP8; MDG = 2.6, *P* < 0.01) at Östergötland, and *Deltaproteobacteria bacterium CG2-30-66-27* (Proteobacteria; MDG = 2.4, *P* < 0.01) at Stockholm. Notable identifications were the differentially abundant bacteria *Sulfurimonas gotlandica* (Proteobacteria; MDG = 0.4, *P* < 0.05) at the Dead Zone (Sörmland) and the archaea *Nitrosopumilus sp. Nsub* (Thaumarchaeota; MDG = 2.2, *P* < 0.01) at the Southern Baltic. Although the alpha diversity of the microbial communities by region differed significantly, no decreasing trend like the ARGs was observed from the northern to the southern regions (**Supplementary Fig. S7A-B**). This was supported by the insignificant correlation of microbial Chao1 richness (*adj. R^2^* = 0.0481; *P* = 0.0522) and Shannon diversity (*adj. R^2^* = −0.0175; *P* = 0.9937) against haversine distance (**Supplementary Fig. S7C-D**). However, PCoA based on Bray-Curtis dissimilarity revealed the differentiation of the microbial communities of the Dead Zone (Sörmland), some stations in the Dead Zone (Mid-South), and the Southern Baltic against the northern regions (**Fig. 2I**). This suggests regional variations in microbial community composition and structure across the Baltic Sea, possibly influenced by local environmental factors and ecological interactions within each region.

A total of 167 MRGs were identified along the Baltic Sea, with the top resistance genes mainly involved with hazardous heavy metals, i.e., *acr3* (4.6% of the total predicted MRG relative abundance), *arsB* (15.2%), and *pstB* (3.8%) conferring resistance to arsenic, *chrA* (4.2%) and *ruvB* (5.3%) to chromium, *actP* (11.7%), *copA* (4.2%) and *copB* (6.7%) to copper (**Supplementary Fig. S4C and Fig. S8A**). Interestingly, the two most abundant predicted MRGs showed regional differentiation, with the arsenic resistance protein (*arsb*) significantly more abundant in the southern regions, except for Östergötland (**Supplementary Fig. S8B**; ANOVA, *df* = 7, *F* = 28.1, *P* < 0.0001). The copper-translocating P-type ATPase (*actP*) was more abundant in the northern areas. However, only the Dead Zone (Sörmland) samples from the southern areas showed a significant difference in *actP* abundance against the northern regions (**Supplementary Fig. S8C**; ANOVA, *df* = 7, *F* = 3.2, *P* < 0.01). To assess the mobility and pathogenic factors potentially involved in disseminating antimicrobial resistance, we also identified the MGEs and VFGs from the dataset. A total of 512 MGE genes were identified and further categorized into functions mediating integration/excision-related genes (57% of the total VFG relative abundance), phage (19%), replication/recombination/repair (15%), stability/transfer/defense (5%), and transfer (3%) (**Supplementary Fig. S4D**). A total of 195 VFGs were identified with the dominant virulence factor types associated with adherence functions (total RPKM relative abundance of 34%), followed by immune modulation (19%), nutritional/metabolic factor (12%), effector delivery system (11%), and stress survival (10%) in all samples (**Supplementary Fig. S4E**).

The trends in alpha diversity observed from the ARGs were relatively similar to the MGEs (**Supplementary Fig. S7E-G**) and the MRGs (**Supplementary Fig. S7H-J**). On the other hand, VFGs showed significantly high Chao1 richness and Shannon diversity in the Dead Zone (Sörmland) region (**Supplementary Fig. S7K-M**). The gene-level composition of the MRGs, MGEs, and VFGs was also visualized using PCoA analysis, showing significant differentiation by regions similar to the ARGs and microbial communities (**Fig. 2K-L**).

### Driving factors of the environmental resistome and co-occurrence patterns

Significant distance decay relationships were reported for the predicted ARG, microbial communities, MRG, MGE, and VFG profiles along the 1,150 km distance across the Baltic Sea (**Fig. 3A**). Testing the linear correlation between Bray-Curtis dissimilarity and haversine distance revealed that compositional dissimilarity increased from north to south, suggesting that microbial communities, and their resistance genes and genetic elements had geographic differences. However, the strengths of the relationships were relatively low, with distance between sites accounting for 2% to 28% of the Bray-Curtis distances. Excluding the hypoxic (< 2 mg/L O_2_) dead zone regions from the model still resulted in relatively low to mild influences accounting for 9% to 45% of the Bray-Curtis distances (**Supplementary Fig. S9**).

**Fig. 3.**
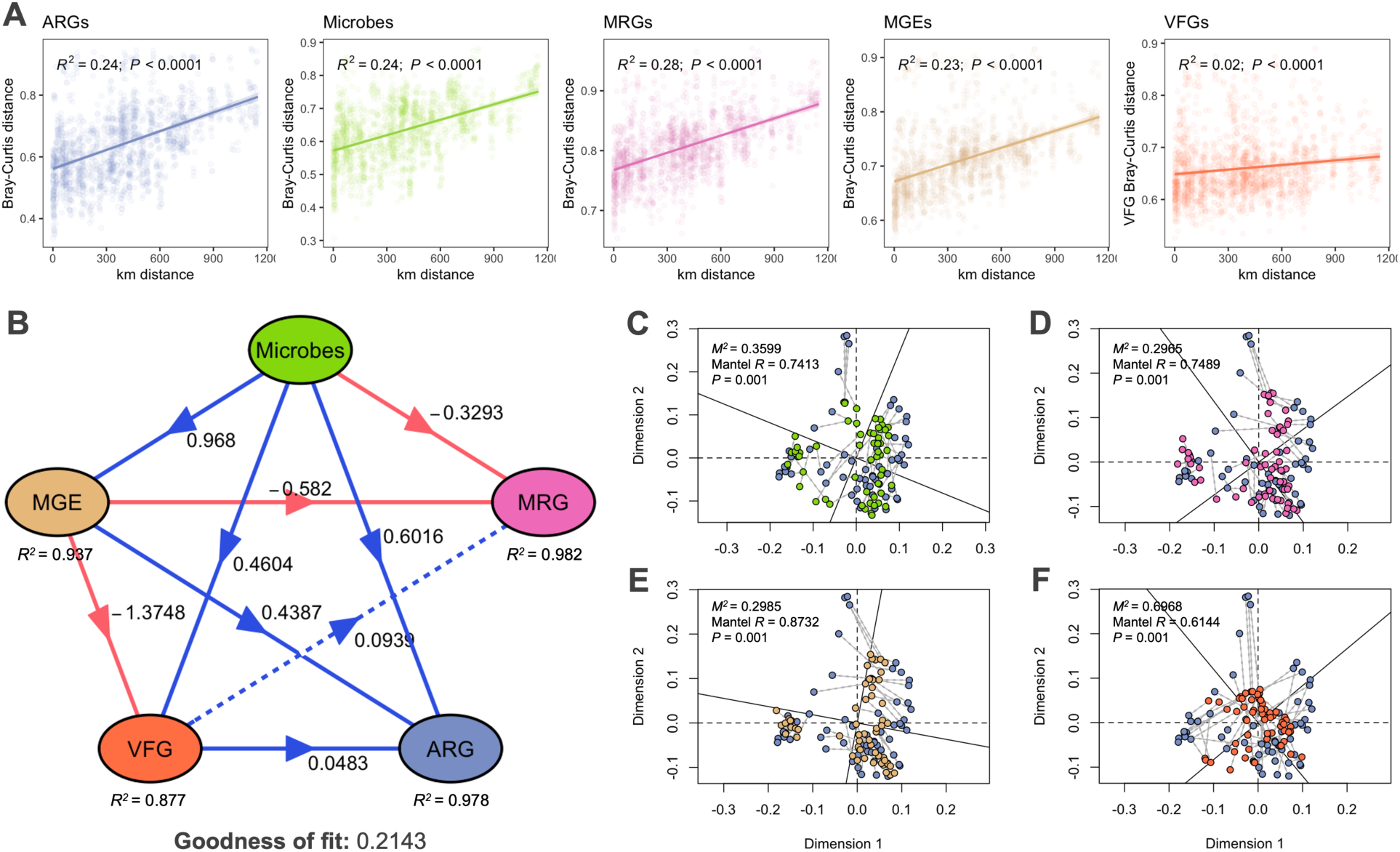
Relationships between resistance and other associated genes. **A** Distance decay relationship of the Bray-Curtis dissimilarities of the predicted ARG, microbial community, MRG, MGE, and VFG profiles with geographic distance (in km). Different colors correspond to the gene profiles in **Fig. 3B**. **B** Specific causality among the microbial community and gene profiles using partial least squares path modeling (PLS-PM). Only the direct effects coefficient is visualized. Total, direct, and indirect effects coefficients are plotted in **Fig. S8**. Arrows represent unidirectional relationships among latent variables. Blue and red lines denote positive and negative relationships, respectively. Dashed lines indicate non-significant coefficients (Pr(>|t|) > 0.05). Procrustes analysis and Mantel test showing the correlation of predicted ARG profile with the **C** microbial communities, **D** MRG, **E** MGE, and **F** VFG profiles.

PLS-PM analysis was performed to quantify the direct and indirect effects of microbial communities and gene profiles in the Baltic Sea (**Fig. 3B**, **Supplementary Fig. S10**, and **Supplementary Table S14**). The model showed that microbial communities, as the exogenous variable, have a substantial direct effect on MGEs (0.97) and a moderate effect on predicted ARGs (0.60). The strong direct effect on MGEs indicates that microbial communities strongly influence the presence of MGEs along the Baltic Sea. We found low direct (0.46) and high negative total effect (−0.87) of microbial communities on VFGs. Additionally, the microbial community has negative direct (−0.33) and strong total (−0.97) effects on MRG composition. The MGE profile significantly negatively affects VFGs (−1.37) and MRGs (−0.58). Meanwhile, VFGs have a minor positive direct effect on predicted ARGs (0.0483) and MRGs (0.0939), with the latter being insignificant at 0.05. However, our model’s relatively low Goodness-of-Fit value of 0.2143 indicates a moderate overall fit. Although the model performed well in predicting specific outcomes, it may not capture all aspects of the relationships between the microbial community and gene profiles. Hence, we further assessed the association between predicted ARG composition, microbial communities, and their genetic elements using Procrustes analyses and Mantel tests (**Supplementary Table S15** and **S16**). The tests revealed a strong and significant correlation between the predicted ARG composition and microbial communities (**Fig. 3C**; *M²* = 0.3599, *P* = 0.001; Mantel *R* = 0.7413, *P* = 0.001), suggesting a closely aligned relationship between these two datasets. Similarly, the predicted ARG composition was significantly correlated with MGEs (**Fig. 3D**; *M²* = 0.2965, *P* = 0.001; Mantel *R* = 0.7489, *P* = 0.001), highlighting the role of microbes and their MGEs in antimicrobial resistance mechanisms. On the other hand, a strong correlation was observed between the predicted ARGs and MRGs (**Fig. 3E**; *M²* = 0.2985, *P* = 0.001; Mantel *R* = 0.8732, *P* = 0.001), which indicates significant co-occurrence and potential co-selection between these resistance mechanisms. Lastly, the correlation between the predicted ARGs and VFGs was weak (**Fig. 3F**; *M²* = 0.6968, *P* = 0.001; Mantel *R* = 0.6144, *P* = 0.001).

### The influence of environmental gradients on the Baltic Sea resistome

Seven environmental factors measured on all 59 monitoring stations were used in the downstream analyses (**Supplementary Table S1** and **Fig. S1**). Water depth ranged from 13 to 125 m. Distinct salinity and temperature gradients from the northern to southern regions were observed, with salinity values ranging from 2.6 to 16.2 and temperature from 0.23 to 11.6°C. Dissolved O_2_ levels ranged from 0.23 to 11.49 mg/L, with the hypoxic Dead Zone regions not having > 0.5 mg/L concentrations. The C/N ratio – a measure of organic matter decomposition varied from 7.61 to 13.81. The δ^13^C and δ^15^N isotope compositions ranged from 2.23‰ to 4.97‰ and from 19.87‰ to 25.75‰, respectively.

Pearson’s correlation between the environmental factors and ARG alpha diversity revealed that salinity, temperature, and δ^13^C had significant relationships with predicted ARG diversity and richness (**Fig. 4A**). In particular, salinity and temperature were negatively correlated with Chao1 richness and Shannon diversity, which suggest that these environmental gradients create conditions that decrease predicted ARG richness and diversity across the Baltic Sea. The southern regions had higher salinity levels that present osmotic stress to most microorganisms, reducing their viability and promoting community composition shifts. This may result in a decrease in ARG-carrying species, as only salt-tolerant microorganisms are selected to survive and dominate in the area. Additionally, elevated temperatures create thermal stress that strongly influences microbial metabolism and growth, which may reduce microbial diversity by eliminating intolerant species. On the other hand, δ^13^C was positively correlated with Chao1 richness and Shannon diversity. GAMs were used to further examine non-linear relationships and assess the multiple interactions of environmental factors on the predicted ARG richness, diversity, and evenness (**Supplementary Table S17**). The model for Chao1 richness indicated that salinity (*edf* = 2.3, *F* = 4.5, *P* < 0.01) and C/N ratio (*edf* = 1, *F* = 4.9, *P* < 0.05) significantly explained 48.8% of the index’s variation, suggesting their importance in shaping the predicted ARG richness. The Shannon diversity model accounts for 54.4% of the variation, with significant contributions from salinity (*edf* = 1.3, *F* = 12.9, *P* < 0.001), C/N ratio (*edf* = 1, *F* = 21.5, *P* < 0.001), and δ^13^C (*edf* = 2, *F* = 6.9, *P* < 0.01) and a marginally significant effect from temperature (*edf* = 2.1, *F* = 3.2, *P* < 0.05). Lastly, the Pielou evenness model showed that depth (*edf* = 1, *F* = 9.1, *P* < 0.01), salinity (*edf* = 1, *F* = 12.5, *P* < 0.01), C/N ratio (*edf* = 7.7, *F* = 5.7, *P* < 0.001), and δ^15^N (*edf* = 2.5, *F* = 3.2, *P* < 0.05) collectively explained 75% of ARG evenness variation.

**Fig. 4.**
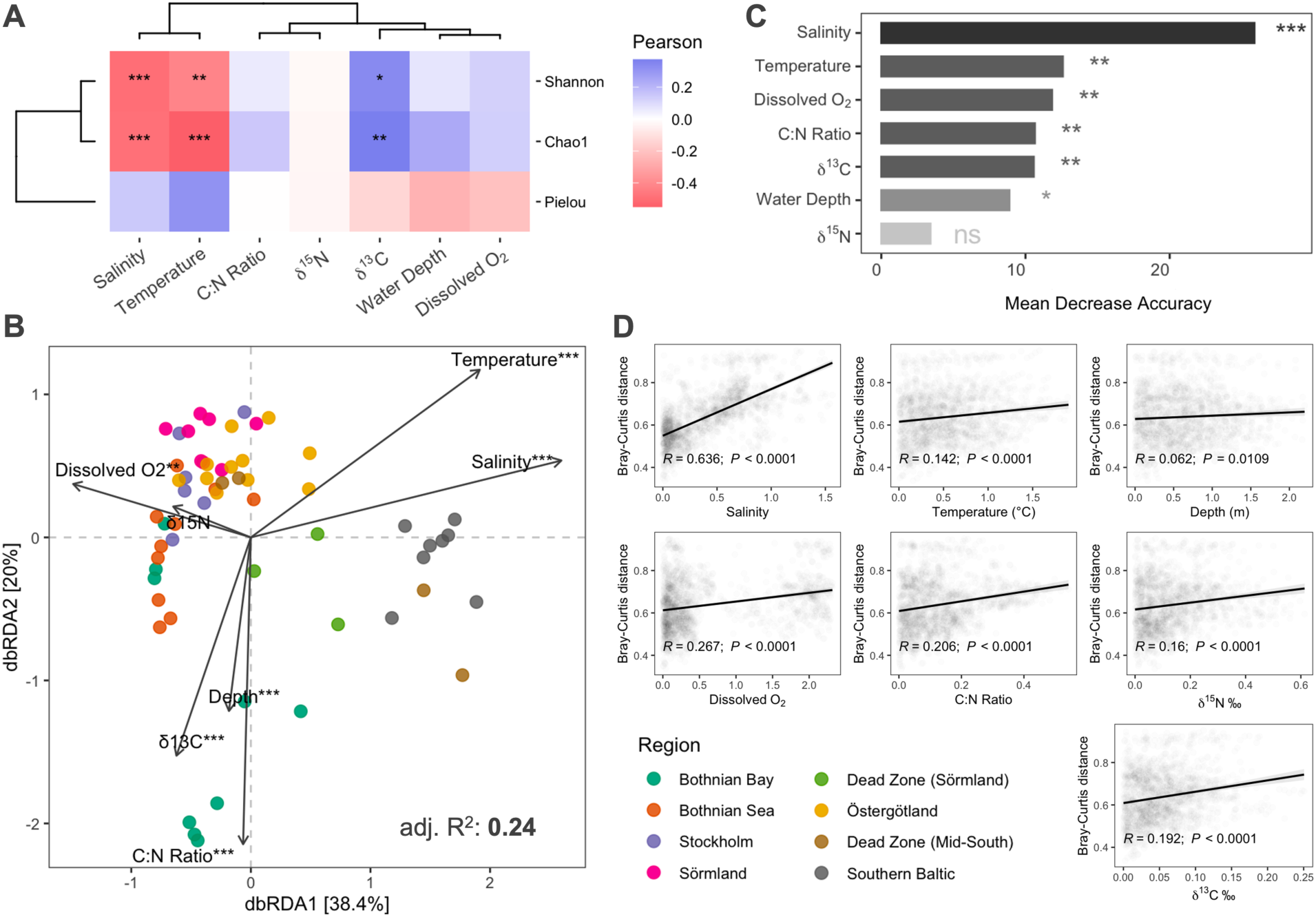
Relationship between the environmental factors and ARG profile. **A** Pearson’s correlation of the relationship between the environmental factors and alpha diversity. The environmental data was log-transformed before the analysis. Positive correlations are indicated in blue, and negative correlations in red. **B** Distance-based redundancy analysis (dbRDA) ordination plot of the predicted ARG composition based on Bray-Curtis distance and the log1p-transformed environmental factors. Significance was obtained by 999 permutations. **C** Random forest analysis based on the contributions of environmental factors on predicted ARG composition. **D** Pearson’s correlation of the relationship between the Euclidean distance of the environmental factors and predicted ARG composition based on Bray-Curtis distance. “***” indicates a significant *P* < 0.001, “**” *P* < 0.01, “*” *P* < 0.05, and “ns” no significant difference.

To further evaluate the relationships between the predicted ARGs and environmental factors, we performed a dbRDA analysis on predicted ARG composition based on Bray-Curtis distance (**Supplementary Table S18**). We found that all the environmental factors assessed, except for the isotope ratio δ^15^N, significantly influenced the regional composition of the predicted ARGs in the Baltic Sea (**Fig. 4B**). In particular, salinity, temperature, C/N ratio, and δ^13^C were the most influential environmental factors affecting the predicted ARG composition in the Baltic Sea. These factors explain a substantial portion of the variability in composition, with salinity (59%) and C/N ratio (76%) being strongly significant. Water depth and dissolved O_2_ also contribute, though to a lesser extent, explaining only 22% and 17% of the variation. Furthermore, we assessed the correlation between these environmental factors and predicted ARG composition. Using a random forest analysis (**Fig. 4C**), salinity was identified as the most important variable influencing ARG composition, having the highest mean decrease accuracy (MDA) value (MDA = 25%, *P* < 0.001). Temperature was the second most important variable (MDA = 14%, *P* < 0.001), followed by dissolved O_2_ (MDA = 12%, *P* < 0.001), C/N ratio (MDA = 12%, *P* < 0.01), δ^13^C (MDA = 11%, *P* < 0.01), and water depth (MDA = 9%, *P* < 0.05). δ^15^N was identified as the least important variable (MDA = 4%, *P* = 0.25), with statistically insignificant importance, suggesting that its effect could be due to random chance. Furthermore, salinity had the strongest correlation with Bray-Curtis distance (R = 0.636, *P* < 0.0001), supporting the aforementioned significant influence of this environmental factor in driving the differences in predicted ARG composition (**Fig. 4D**). Other environmental factors, i.e., dissolved O_2_ (*R* = 0.267, P < 0.0001), C/N ratio (*R* = 0.206, *P* < 0.0001), δ^15^N (*R* = 0.16, P < 0.0001), δ^13^C (*R* = 0.192, P < 0.0001), and temperature (*R* = 0.142, *P* < 0.0001), showed weak but still significant correlations. In contrast, water depth (*R* = 0.062, *P* = 0.0109) had the weakest correlation among the factors analyzed.

Correlations between ARG types and environmental factors were performed to present the complex interactions between the predicted ARG composition at a higher classification level and the Baltic Sea environmental gradient (**Fig. 5** and **Supplementary Table S19**). Hierarchical clustering was used to further classify the relationships among ARG types and environmental factors based on their correlation patterns. Salinity showed strong positive correlations with several ARG types, e.g., glycopeptide, aminoglycoside, diaminopyrimidine, polymyxin, and pleuromutilin, including the unclassified ARGs. Temperature had the most significant positive correlations with mostly dominant ARG types, e.g., multidrug, glycopeptide, beta-lactam, bacitracin, aminoglycoside, and tetracycline resistance, which indicates that higher temperatures may create favorable conditions for the proliferation of the hosts carrying these resistance genes.

**Fig. 5.**
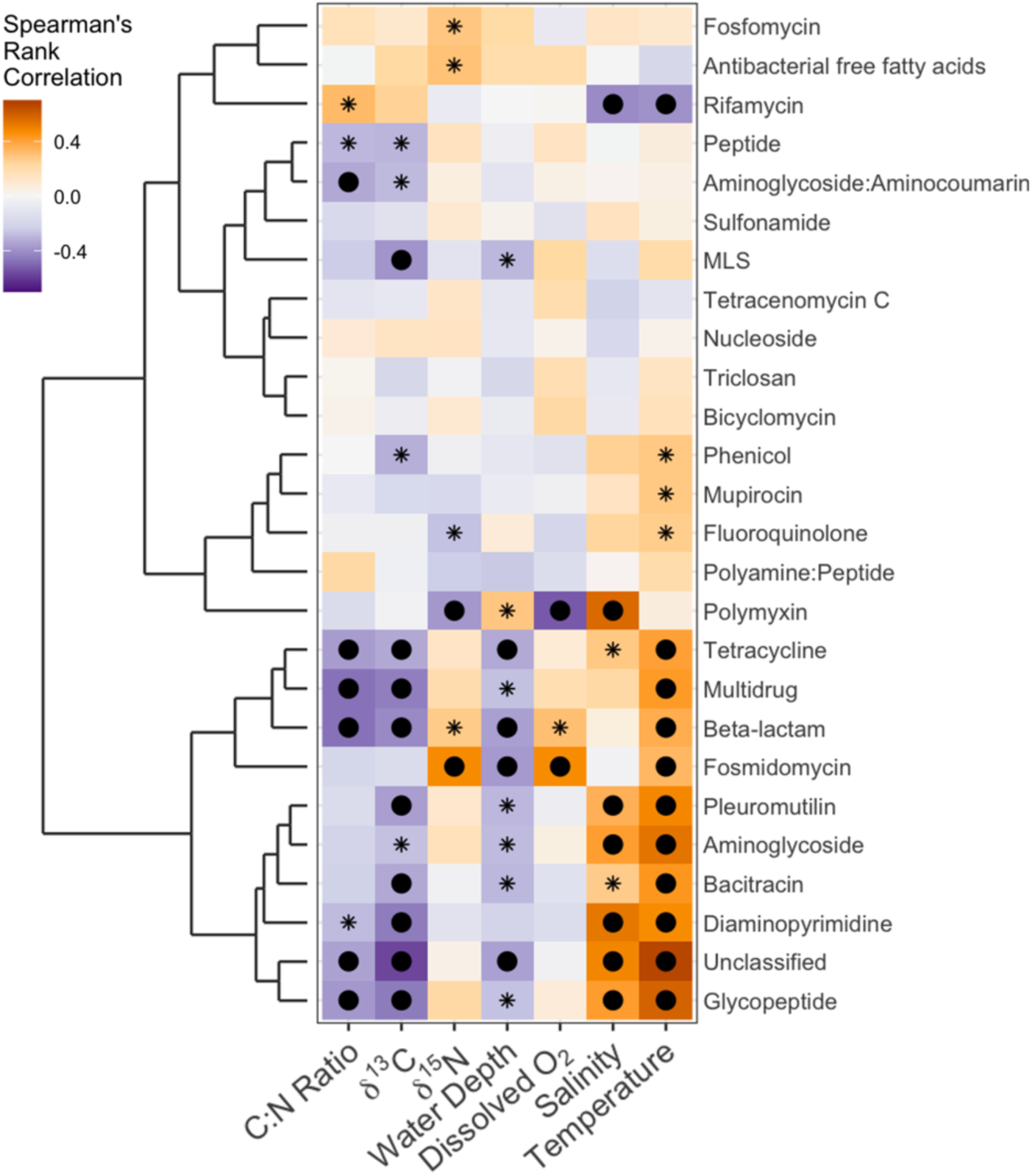
Spearman’s rank correlation on the relationship between the environmental factors and ARG types. Positive correlations are indicated in yellow, and negative correlations in purple. The asterisk indicates p < 0.05, and the filled circle indicates FDR corrected *P* < 0.05.

## Discussion

Understanding the relationships between environmental factors, microbial communities, and their resistance mechanisms can provide knowledge on the dynamics of environmental resistomes in aquatic ecosystems that pose growing public health concerns [4, 56]. This includes exploring the main drivers and mechanisms underlying resistance and its associated genes and their evolution, selection and spread in natural environments [57, 58], and the knowledge of the background levels of antimicrobial resistance in potential natural reservoirs [3]. Using the metagenomics data from sediments sampled across the Baltic Sea [27], we characterized the environmental resistome of its benthic ecosystem over spatial distances, specifically focusing on resistance, i.e., ARGs and MRGs, and their mobility and pathogenicity genes.

### Salinity and temperature modulated the distribution patterns of the resistome

Salinity and temperature have been identified as key drivers of ecosystem dynamics, biodiversity, and biogeochemical processes across the Baltic Sea [26, 27, 59, 60]. Its pronounced environmental gradient is characterized by fresher and cooler northern waters compared to the south’s relatively saline and warmer conditions [61]. We found that the environmental resistome also strongly correlates with the salinity gradient, with the predicted ARG diversity declining with increasing salinity along the 1,150 km of the Baltic Sea, and its composition significantly differentiated across regions. These results are in accordance with studies that reported salinity as a key factor in ARG composition in aquatic ecosystems [15, 62–65]. Salinity is a crucial determinant of microbial community composition, mainly by increasing the selective pressure on microorganisms [66], which might consequently influence the proliferation of ARGs [67, 68]. Tee et al. [69] observed that similarities in the community composition of planktonic and benthic microorganisms decline as the differences in salinity and geographic distance increase. Accordingly, microorganisms exhibited varying tolerances to salinity, resulting in distinct populations across the Baltic Sea [70]. In this study, we observed that the microbial community and the predicted regional ARG composition with higher salinity significantly differed from areas with lower to mid-saline conditions. However, we only observed significant differences in the richness and diversity of the predicted ARGs and not with the microorganisms, which suggests that ARG distribution is more influenced by microbial community composition than species diversity. Likewise, Ohore et al. [65] demonstrated that the salinity gradient between freshwater and estuary environments had more influence on pelagic bacterial composition than species diversity. This indicates that the composition of the resident species thriving in more saline environments influenced the ARG profile more than local species diversity.

We also observed that the temperature gradient might have played a role in structuring ARG distribution. Warmer temperatures in the Southern Baltic may influence microbial growth and metabolic activities, directly influencing the proliferation of specific ARGs in the region. Yu et al. [71] reported that in elevated water temperatures, ARG diversity was reduced while its relative abundance increased alongside several opportunistic pathogens. Temperature may influence the permeability of cell membranes and inactivate metabolic enzyme activities involved in resistance mechanisms, with higher temperatures also resulting in the inability of sensitive species to survive [72]. Accordingly, we observed similar patterns with the richness and diversity of our predicted ARGs having significant negative correlations with temperature. The combined effects of temperature and salinity may have a concurrent influence on microbial community structure and biochemical composition [73]. The combined stress of increasing temperature and salinity could synergistically enhance the selection of microorganisms that are both thermotolerant and halotolerant, potentially concentrating ARGs within these adapted taxa at specific regions.

We also observed that the C/N ratio has influenced a substantial portion of the variability in ARG composition. The C/N ratio indicates the relative amounts of carbon and nitrogen in the environment and is a proxy of organic matter quality and resource availability for benthic microbial communities. It is an important factor in shaping the composition of local communities through its influence on metabolic processes and community dynamics [e.g., 74]. Although we did not observe a significant influence of the C/N ratio on predicted ARG diversity, we identified that δ^13^C was significantly positively correlated with richness and diversity. This stable isotopic measurement was used to trace carbon sources to understand environmental carbon cycling [75]. The positive correlation indicates that Baltic Sea regions with higher δ^13^C values, which typically reflect greater marine-derived carbon inputs and higher primary productivity [76], also had higher diversity. This relationship suggests that an increased proportion of marine organic matter or higher productivity supports more complex and varied microbial communities, which would most likely harbor highly diverse ARGs. High organic carbon concentrations would enhance microbial growth and metabolic activity and promote horizontal gene transfer and, consequently, the proliferation of ARGs in the environment [77].

In principle, the transmission, persistence, and enrichment of ARGs in natural ecosystems are ecological processes that are affected by multiple factors [78]. We found that the combined effects of salinity and temperature gradients, alongside carbon availability, create a complex environmental landscape that shapes the diversity and distribution of ARGs in the Baltic Sea. Understanding the relationship between these environmental factors, specifically salinity, and ARG diversity and composition is crucial for managing the antimicrobial resistance in marine ecosystems, particularly in regions like the Baltic Sea, where salinity levels vary. Also, identifying salinity and temperature as key factors in the Baltic benthic ecosystem highlights the importance of further research into how these environmental factors influence the environmental resistome, which could inform strategies for the emergence and dissemination of antimicrobial resistance in complex aquatic ecosystems in a changing environment.

### Biogeographic pattern of the environmental resistome across the Baltic Sea

From the 59 benthic metagenomes, we predicted 378 ARGs conferring resistance to 26 antibiotic classes (i.e., ARG types). Likewise, profiling of the global ocean resistome reported genes conferring putative resistance to 26 antibiotic classes [79], while another global assessment of marine environments identified 23 types [80]. Predicted genes associated with multidrug resistance were the most dominant ARG type, with higher prevalence and abundance in the northern Baltic regions. At the gene level, the multidrug ATP-binding cassette (ABC) transporter gene was the most dominant predicted gene conferring multidrug resistance. This gene encodes for a widely distributed and functionally diverse membrane protein responsible for ATP-driven substrate translocation. It is a crucial multidrug resistance mechanism as it encodes proteins that actively pump antimicrobials out of cells, reducing their efficacy [81]. Glycopeptide resistance was the second most dominant ARG type, with increasing prevalence and abundance in the southern areas. From this ARG type, the glycopeptide resistance-associated *vanR* gene was the most dominant. This is an OmpR-family transcriptional activator in the VanSR regulatory system, which controls how the expression of vancomycin resistance is induced [82]. Both multidrug and glycopeptide resistance types were identified as the most dominant ARG types from the core marine resistome of globally distributed locations [80]. They were also reported as the most ubiquitous ARG types in resistant bacteria from urban environments [83].

We report spatial variation in the environmental resistome of the Baltic Sea, with higher diversity observed in the northern regions, i.e., Bothnian Bay, Bothnian Sea, Stockholm, and Sörmland, and lower diversity for the dead zones and the southern areas. The predicted ARG diversity and composition we observed from the northern regions suggest that the resistome and microbial community profiles in these areas were complex and capable of supporting a wide range of resistance mechanisms. In contrast, the dead zones and the Southern Baltic Sea showed lower predicted ARG diversity, indicating significantly varied within-group composition, with a relative decrease in predicted multi-drug resistance and increased abundance of predicted genes associated to glycopeptide resistance and other types. This might result from the region’s different environmental conditions and selective pressures, leading to specific microbial communities associated with these resistance types. In particular, the dead zones are characterized by low O_2_ levels due to eutrophication. These hypoxic conditions govern the microbial community assembly [84], favoring anaerobic or facultative anaerobic species with specific ARGs different from those in more oxygen-rich environments [85]. Interestingly, the most dominant MRG, the arsenic resistance protein (*arsb*), was significantly abundant in the southern regions, except for the Östergötland samples. Arsenic-based Chemical Warfare Agents (CWA) are anthropogenic sources of arsenic in the bottom sediments of the Baltic Sea [86], with concentrations ranging from less than 5 μg g^−1^ up to 29 μg g^−1^ in the sediments of the southern region. These concentrations are considered low and do not pose serious threats to marine organisms [87]. However, localized hotspots of arsenic contamination could possibly have led to the evolution and proliferation of arsenic resistance mechanisms in the exposed benthic microbial communities of the area.

The decreasing predicted ARG richness and diversity from north to south reflects the varying environmental pressure across the regions. The geographic differences in antimicrobial resistance patterns highlight the interplay between environmental factors, potential anthropogenic pressure, and microbial dynamics, emphasizing the importance of considering various environmental gradients when assessing ecosystem health and resilience for managing antimicrobial resistance in potential natural reservoirs of antimicrobial resistance, e.g. the Baltic Sea ecosystem. Previous studies reported that anthropogenic pollution enhanced the relative abundance of dominant ARG hosts and increased the co-occurrence of multiple ARGs from pristine to urban rivers [88]. Accordingly, Provencher et al. [89] profiled ARGs in the Canadian High Arctic and observed similar findings where areas with higher anthropogenic impact had greater ARG diversity than less impacted areas. Environmental stressors can further exacerbate these effects by disrupting microbial communities, favoring more resilient and resistant organisms, which might lead to ecosystem imbalances, affecting critical functions, e.g., nutrient cycling, and promoting the spread of resistance across ecosystems.

### Microbial communities and mobile genetic elements were significantly correlated with the environmental resistome

Microbial communities and functional genes played a significant role in establishing resistome profiles in the environment [90]. As expected, we found that benthic microbial communities across the Baltic Sea had strong direct effects on the MGE profile but had intermediate direct effects on predicted ARG composition. Microbial communities play crucial roles in determining the presence and distribution of MGEs, which are key factors for gene transfer, including ARGs [91]. Previous studies reported strong direct effects of MGEs on ARG composition where horizontal gene transfer-mediated MGEs were the major mechanisms involved in disseminating ARGs in aquatic ecosystems [e.g., 92-95]. Consequently, while the direct effect of microbial communities on our predicted ARG composition was moderate, they exerted strong indirect effects. This suggests that microbial communities influence other factors, i.e., MGEs, which mainly contribute to structuring ARG composition in the environment [96]. Previous studies have provided evidence on the influence of microbial community composition and species diversity on ARG dynamics [e.g., 97, 98]. Microbial community composition directly impacts intrinsic ARG prevalence by influencing the presence of specific resistant species [99] and the potential for horizontal gene transfer [100]. Additionally, species diversity may influence ARG dynamics through ecological interactions. Higher species diversity can also limit the dominance of antimicrobial-resistant species through competition, reducing resistance levels [101, 102], but may also promote the co-occurrence and transfer of ARGs between species [103].

We also found strong positive correlations between the predicted ARG and MRG composition. These two resistance genes were reported to have significant correlations in aquatic systems [20], which suggests that the mechanisms driving resistance to antimicrobial agents and heavy metals might be interconnected. However, these observations were uncommon in environmental bacterial communities and require further evaluation [104]. Interestingly, although we found strong correlations between the two resistance gene profiles, our model revealed negative direct effects of microbial communities on MRG composition, which suggests that as the diversity and composition of microbial communities increased, the prevalence of MRGs decreased. This also indicates that limiting ecological interactions influences the spread of metal resistance within diverse microbial communities.

Furthermore, we observed that the benthic microbial communities had a negative total effect on VFGs, indicating a potential competitive interaction where more diverse microbial communities overturn virulence factors. This suggests that changes in microbial composition may be associated with a reduction in VFGs, which could indicate competitive interactions where more diverse microbial communities suppress pathogenicity factors. The predicted ARGs and VFGs also had weak correlations, which indicates that the factors influencing the distribution and abundance of ARGs differ from those affecting host pathogenicity and vice versa. This contrasts with previous studies that reported positive correlations between ARGs and VFGs [105, 106]. While both are important for microbial survival and pathogenicity, they may be regulated by distinct mechanisms and respond differently to environmental factors. Hence, virulence factors may be more influenced by host-pathogen interactions [107], which might explain its weak correlation with the ARG profile. Still, these studies were based on wastewater-impacted and gut microbiome samples which might also explain their observed positive correlations. Further understanding of the independent regulation of ARGs and VFGs can provide valuable insights into the complex dynamics of antimicrobial resistance and pathogenicity in aquatic ecosystems. These findings highlight the complex interactions between microbial communities, resistance genes, and associated functional genes in the Baltic Sea, which are useful for understanding antimicrobial resistance and ecological dynamics in the region.

### Climate change implications on the environmental resistome of the Baltic Sea

The spatial variations in the environmental resistome of the benthic ecosystem and the influence of environmental factors on antimicrobial resistance have significant implications for climate change dynamics in aquatic ecosystems. Climate change is expected to exacerbate existing environmental stressors, altering the biological, chemical, and physical factors in aquatic ecosystems [108]. These stressors have been identified as highly important predictors of antimicrobial resistance risk [58, 109]. For example, climate change-induced shifts in habitat conditions, e.g., salinity and O_2_ levels, can further influence microbial community composition and resistance gene dynamics. Warming temperatures may lead to changes in water stratification and nutrient cycling, affecting the availability of resources for microbial growth and metabolism [110]. These changes may, in turn, impact the distribution and abundance of resistant microorganisms and resistance genes in the Baltic Sea ecosystem.

Understanding the complex interplay between climate change, environmental factors, and antimicrobial resistance dynamics is crucial for developing effective surveillance strategies and ensuring the sustainability of aquatic ecosystems in the face of climate change. In this study, we only assessed spatial dynamics and recommend that future research should investigate the seasonal variations in resistance patterns, given that distinct environmental factors, i.e., temperature and salinity, influence the microbial communities and resistome dynamics in the Baltic Sea. Additionally, comparative studies in similar ecosystems, e.g., the North Sea, could determine if our observed dynamics in the Baltic Sea’s benthic ecosystem are representative of broader trends in antimicrobial resistance across freshwater and brackish environments. Finally, further assessment of the impact of anthropogenic activities, e.g., land use and urbanization around the Baltic Sea, may clarify specific sources of contaminants, enabling targeted monitoring and mitigation based on regional practices.

### Monitoring aquatic ecosystems as potential natural reservoirs of antimicrobial resistance

Our study underscores the significant implications of profiling antimicrobial resistance in aquatic ecosystems and highlights the crucial role of environmental factors in shaping ARG diversity and composition. In aquatic ecosystems influenced by anthropogenic activities and diverse environmental conditions, multiple sources of contamination can be introduced, and the risk of antimicrobial resistance persistence and spread is significantly elevated [20, 88, 89]. Therefore, it is essential to identify and monitor these environments, as they may play a role in the dissemination of antimicrobial resistance [3, 12, 13]. This, alongside the development of monitoring frameworks that focus on integrating the identification of antimicrobial-resistant microorganisms, their resistance mechanisms and functional genes, and mapping contaminant sources, would be crucial for guiding investment in strategies to mitigate the emergence and further spread of resistance between ecosystems [111]. Additionally, the prioritization and enhanced surveillance of potentially high-risk ecosystems could serve as an early warning system for emerging resistance threats, which would be useful in protecting both the environmental and public health sectors [112]. Lastly, long-term studies tracking resistance over time would help detect trends, assess the impacts of mitigation efforts, and support adaptive monitoring approaches. Together, these efforts would advance strategies for monitoring antimicrobial resistance in vulnerable aquatic ecosystems worldwide.

## Conclusion

We presented the environmental resistome, i.e., ARGs and MRGs, of the Baltic Sea benthic ecosystem alongside its microbial communities and specific functional genes, i.e., MGEs and VFGs. We highlight the complex relationships between environmental factors and antimicrobial resistance and identified salinity and temperature as the main environmental drivers influencing the diversity and distribution of ARGs across different geographic regions. We demonstrated that the combined effects of salinity and temperature, alongside nutrient availability, created a complex environmental landscape that modulated the diversity and distribution of ARGs in each Baltic Sea region. We also report the relationships between microbial communities, resistance, and associated functional genes in the benthic ecosystem and emphasize the important role of microbial and MGE composition in ARG distribution. Understanding these dynamics is crucial for managing antimicrobial resistance in aquatic ecosystems, particularly in regions like the Baltic Sea with varying environmental conditions. We recommend that further research into the mechanisms underlying these relationships is needed to inform effective strategies in monitoring the emergence and dissemination of antimicrobial resistance in complex aquatic ecosystems.

## List of abbreviations

ANOVA: Analysis of variance
ARG: Antimicrobial resistance gene
AMR: Antimicrobial resistance
dbRDA: distance-based redundancy analysis
GAM: Generalized additive model
MGE: Mobile genetic element
MLS: Macrolide−lincosamide−streptogramin
mOTU: Metagenomic Operational Taxonomic Units
MRG: Metal resistance gene
PCoA: principal coordinate analysis
TC: Total Carbon
TN: Total nitrogen
RPKM: Reads per kilobase million
VFG: Virulence factor gene.

## Supplementary Information

**Additional file 1** (Supplementary-Information.pdf): **Supplementary Fig. S1** Log-transformed values of the environmental factors visualized per region. Different letters indicate differences between groups tested by ANOVA (p-value <0.05). **Supplementary Fig. S2** Comparison of ARG database and methods. Bar plots of **A** the number of annotations per method/database, **B** ADONIS R^2^ based on the relative influence of Region on the community dissimilarity between methods, and **C** the adjusted R^2^ of the distance-based redundancy analyses (dbRDA). The *** in the adonis results indicate significance at p-value <0.001, and ** at p-value <0.01. **Supplementary Fig. S3** Autocorrelations among the environmental factors. **Supplementary Fig. S4** Relative abundance plots of **A** ARG types, **B** microbial phyla, **C** MRGs, **D** MGE, and **E** VFG categories. **Supplementary Fig. S5 A** Relative abundance boxplots of the top ARG types across regions. Separate boxplot visualizations of the multidrug and glycopeptide resistance types in **Fig. 1D**. **B** Differential abundance test by Kruskal-Wallis rank sum and random forest. The variable importance plot shows the top 15 differentially abundant ARG subtypes with high mean decrease Gini scores. **Supplementary Fig. S6** Differential abundance test by Kruskal-Wallis rank sum and random forest. The variable importance plot shows the 47 differentially abundant species. **Supplementary Fig. S7** Alpha diversity of the **A-B** microbial communities, **C-E** MRG, **F-H** MGE, and **I-K** VFG. Different letters indicate differences between groups tested by ANOVA or Kruskal-Wallis test (p-value < 0.05). **Supplementary Fig. S8 A** Relative abundance boxplots of the top MRG types across regions. Separate boxplot visualizations of the **B** arsenic resistance protein ArsB and **C** copper-translocating P-type ATPase. Different letters indicate differences between groups tested by ANOVA (P-value < 0.05). **Supplementary Fig. S9** Distance decay relationship of the Bray-Curtis dissimilarities of the ARG, microbial community, MRG, MGE, and VFG profiles with haversine distance (in km) in the Baltic Sea without the dead zones samples. **Supplementary Fig. S10** Effects coefficients of the PLS-PM model of the microbial community, ARG, MRG, MGE, and VFG profiles as latent variables.

**Additional file 2** (Supplementary-Tables.xls): **Supplementary Table S1** Metadata: Sample information and environmental factors from Broman et al. (2022). Metagenomics sequence codes, site descriptors, i.e. region, station, haversine distance (km), coordinates, and physicochemical factors, i.e., water depth (m), salinity, temperature, dissolved O2 (mg/L), concentrations of total carbon (TC) and total nitrogen (TN) (µmol/g), and their ratio – C/N, and isotopic signatures, i.e., stable 13C and 15N isotope compositions (‰). **Supplementary Table S2** Metagenomics data processing information: Sequence codes, raw read counts, quality-filtered and adapter trimmed, contaminant filtered, error corrected read counts, and contig assembly stat values. **Supplementary Table S3** Relative abundance table of the microbial community based on mOTUs profiler. **Supplementary Table S4** Comparison of methods for antimicrobial resistance gene (ARG) profiling. **Supplementary Table S5** Antimicrobial resistance gene (ARG) data: Reads per kilobase million (RPKM) absolute counts. **Supplementary Table S6** Mobile resistance gene (MRG) data: RPKM absolute counts. **Supplementary Table S7** Mobile genetic elements (MGE) data: RPKM absolute counts. **Supplementary Table S8** Virulence factor gene (VFG) data: RPKM absolute counts. **Supplementary Table S9** Differential abundance test by Random Forests for ARG subtypes per region. **Supplementary Table S10** Antimicrobial resistance genes (ARG) alpha diversity estimates per sample. **Supplementary Table S11** Statistical results of Levene’s test for homogeneity of variance of the alpha diversity estimates. **Supplementary Table S12** Statistical results of the permutation test for homogeneity of multivariate dispersions and permutation test for adonis under a reduced model for the microbial composition, ARG, MRG, MGE, and VFG data. **Supplementary Table S13** Differential abundance test by Random Forests for the species per region. **Supplementary Table S14** Summary statistics of the partial least squares path model (PLS-PM) in R of the latent variables - microbial communities, MGEs, VFGs, ARGs, and MRGs. **Supplementary Table S15** Statistical results of the Procrustes analysis in R of ARG composition based on Bray-Curtis distance against the microbial communities, MRGs, MGEs, and VFGs. **Supplementary Table S16** Statistical results of the Mantel test in R of ARG composition based on Bray-Curtis distance against the microbial communities, MRGs, MGEs, and VFGs. **Supplementary Table S17** Statistical results of the General additive models (GAMs) in R of the alpha diversity and environmental factors. **Supplementary Table S18** Statistical results of the permutation test for dbRDA analyses in R of the ARG composition and environmental factors. **Supplementary Table S19** Spearman’s rank correlation between ARG alpha diversity metrics and environmental factors.

## Declarations

### Ethics approval and consent to participate

Not applicable.

### Consent for publication

Not applicable.

### Availability of data and materials

All raw sequence data are deposited and available online at the European Nucleotide Archive with an accession number PRJEB41834 (Run accession: ERR5010645-661 and ERR5010683-724). Additional data from the analyses presented in this paper are available in the Supplementary Material, and the corresponding input data and R code used for visualization and statistical analysis are available at https://github.com/jserrana/baltic-amr.

### Authors’ contributions

JMS conceptualized the study, analyzed the data, and drafted the manuscript. All authors read, revised, and approved the final version of the manuscript.

## Acknowledgments

The authors acknowledge the National Academic Infrastructure for Super-computing in Sweden for granting us access and storage to the Uppsala Multidisciplinary Center for Advanced Computational Science (UPPMAX) computational infrastructure.

