## Supplementary Figures for "Environmental drivers of the resistome across the Baltic Sea"

*Authors & Affiliations*

Joeselle M. Serrana[1,2]*, Francisco J. A. Nascimento[1,3], Benoît Dessirier[1,4], Elias Broman[1,3,4], and Malte Posselt[1,2]

[1]Stockholm University Center for Circular and Sustainable Systems (SUCCeSS), Stockholm University, 106 91 Stockholm, Sweden

[2]Department of Environmental Science (ACES), Stockholm University, 106 91 Stockholm, Sweden

[3]Department of Ecology, Environment, and Plant Sciences (DEEP), Stockholm University, 106 91 Stockholm, Sweden

[4]Baltic Sea Centre, Stockholm University, Stockholm, Sweden

*Supplementary Figures*

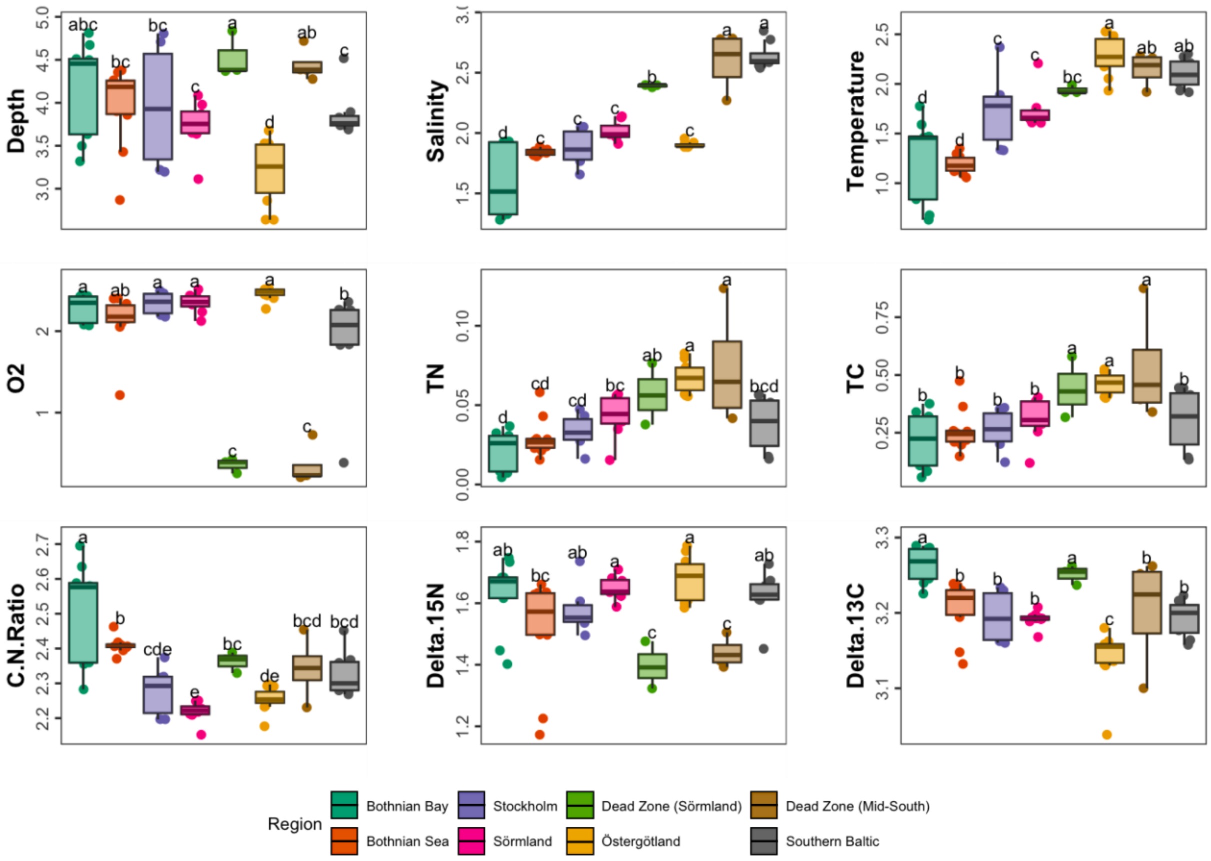


**Supplementary Fig. S1** Log-transformed values of the environmental factors visualized per region. Different letters indicate differences between groups tested by ANOVA (p-value <0.05).

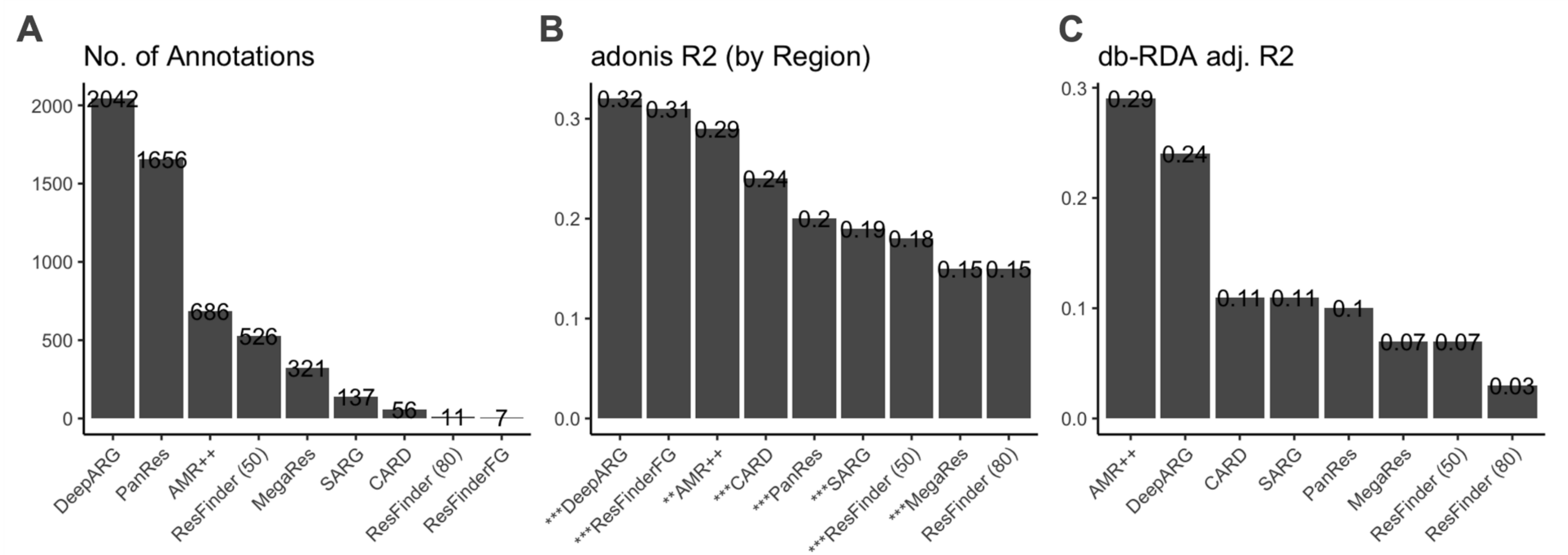


**Supplementary Fig. S2** Comparison of ARG database and methods. Bar plots of **A** the number of annotations per method/database, **B** ADONIS $R^2$ based on the relative influence of Region on the community dissimilarity between methods, and **C** the adjusted $R^2$ of the distance-based redundancy analyses (db-RDA). The *** in the adonis results indicate significance at p-value <0.001, and ** at p-value <0.01.

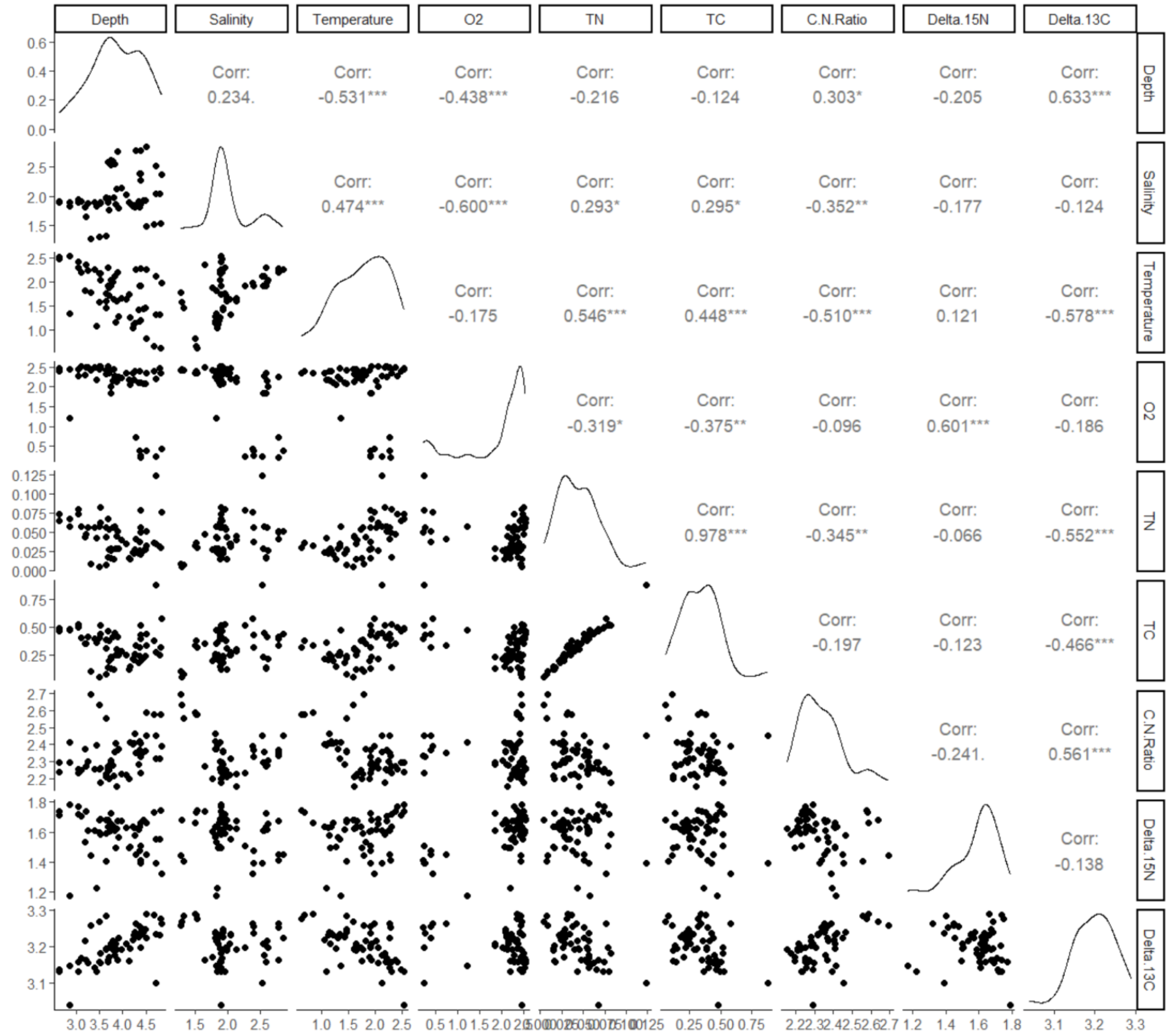


**Supplementary Fig. S3** Autocorrelations among the environmental factors.

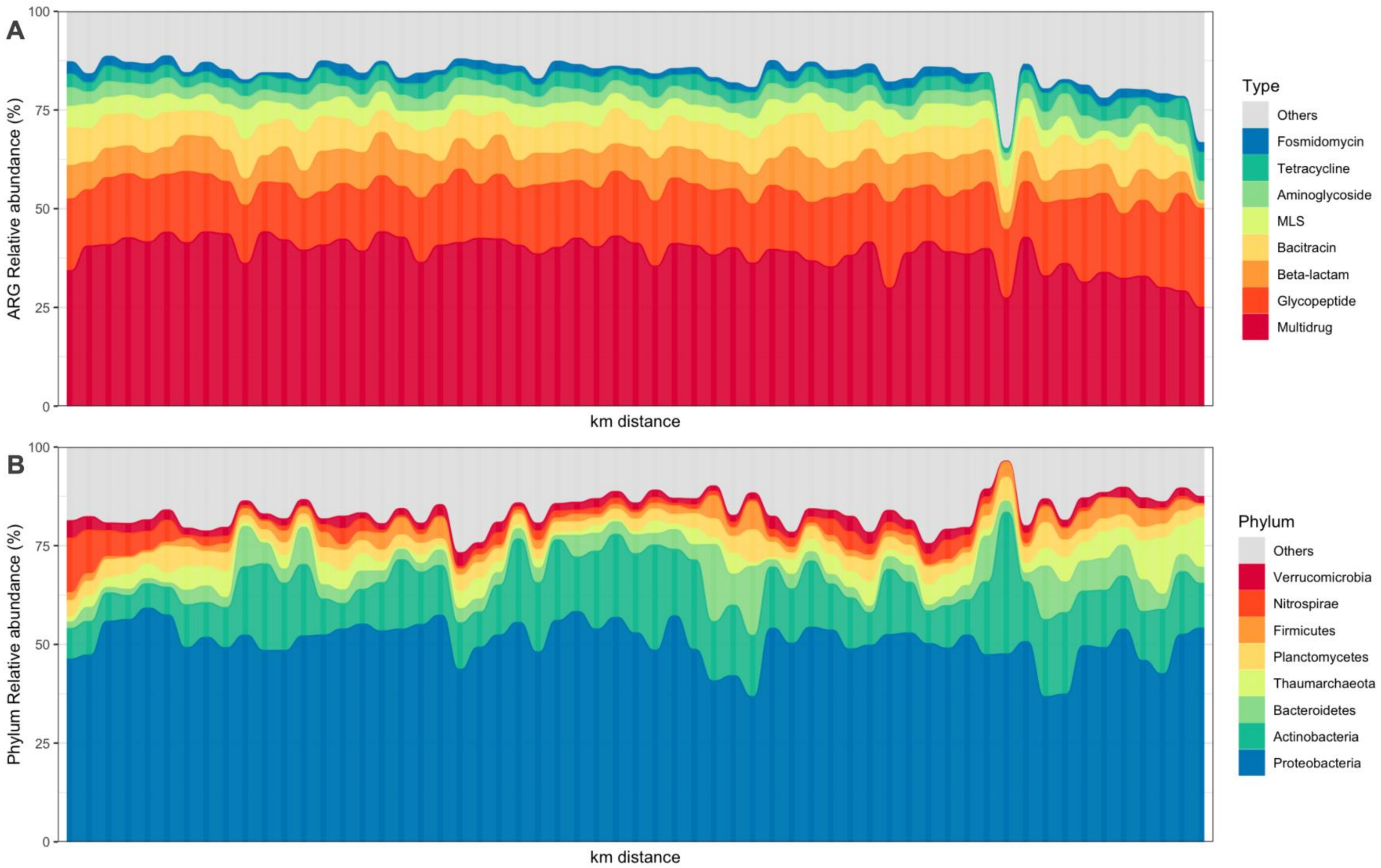


**Supplementary Fig. S4** Relative abundance plots of **A** ARG types, **B** microbial phyla, **C** MRGs, **D** MGE, and **E** VFG categories.

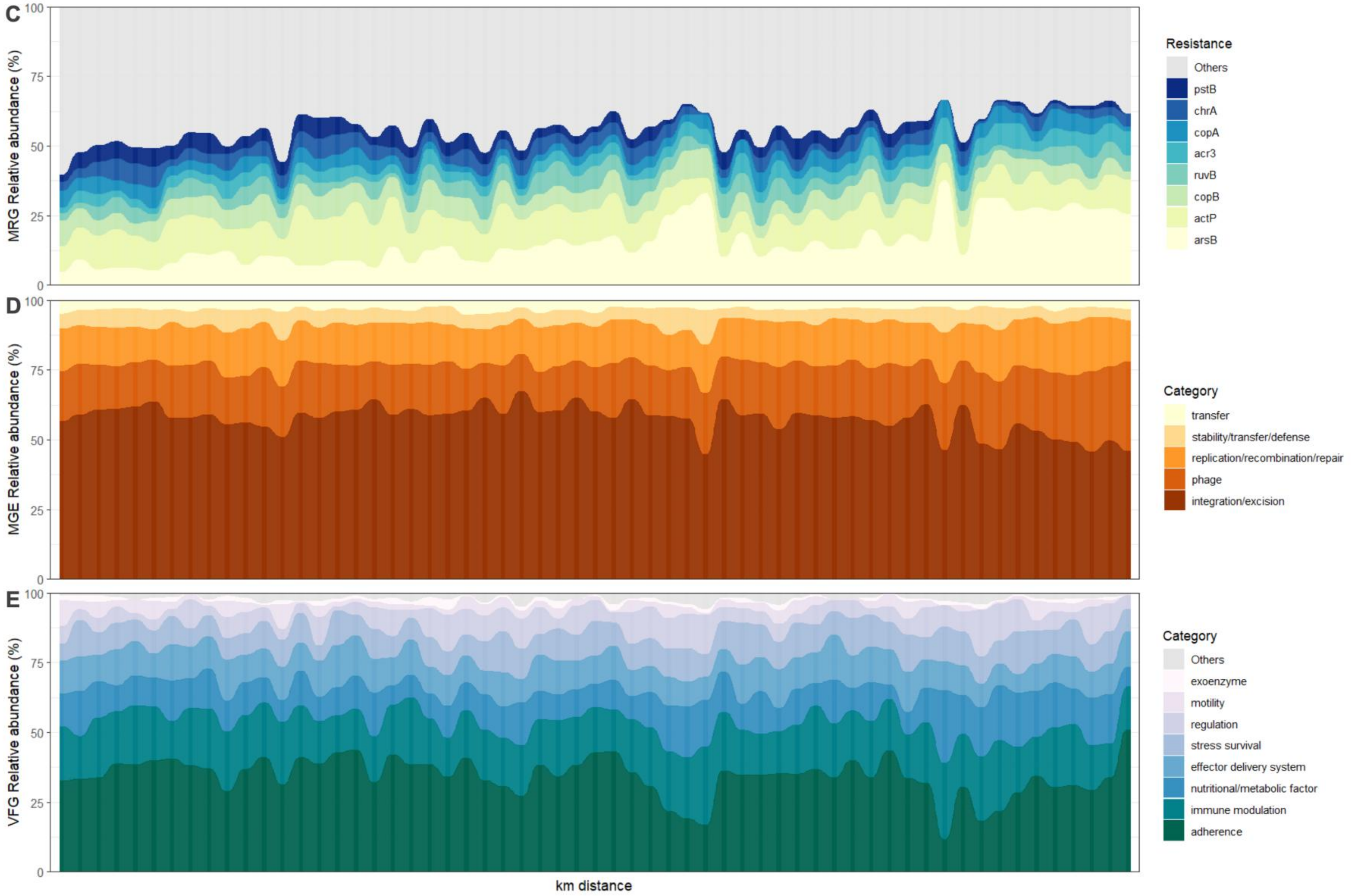


**Supplementary Fig. S4 (cont.)** Relative abundance plots of **A** ARG types, **B** microbial phyla, **C** MRGs, **D** MGE, and **E** VFG categories.

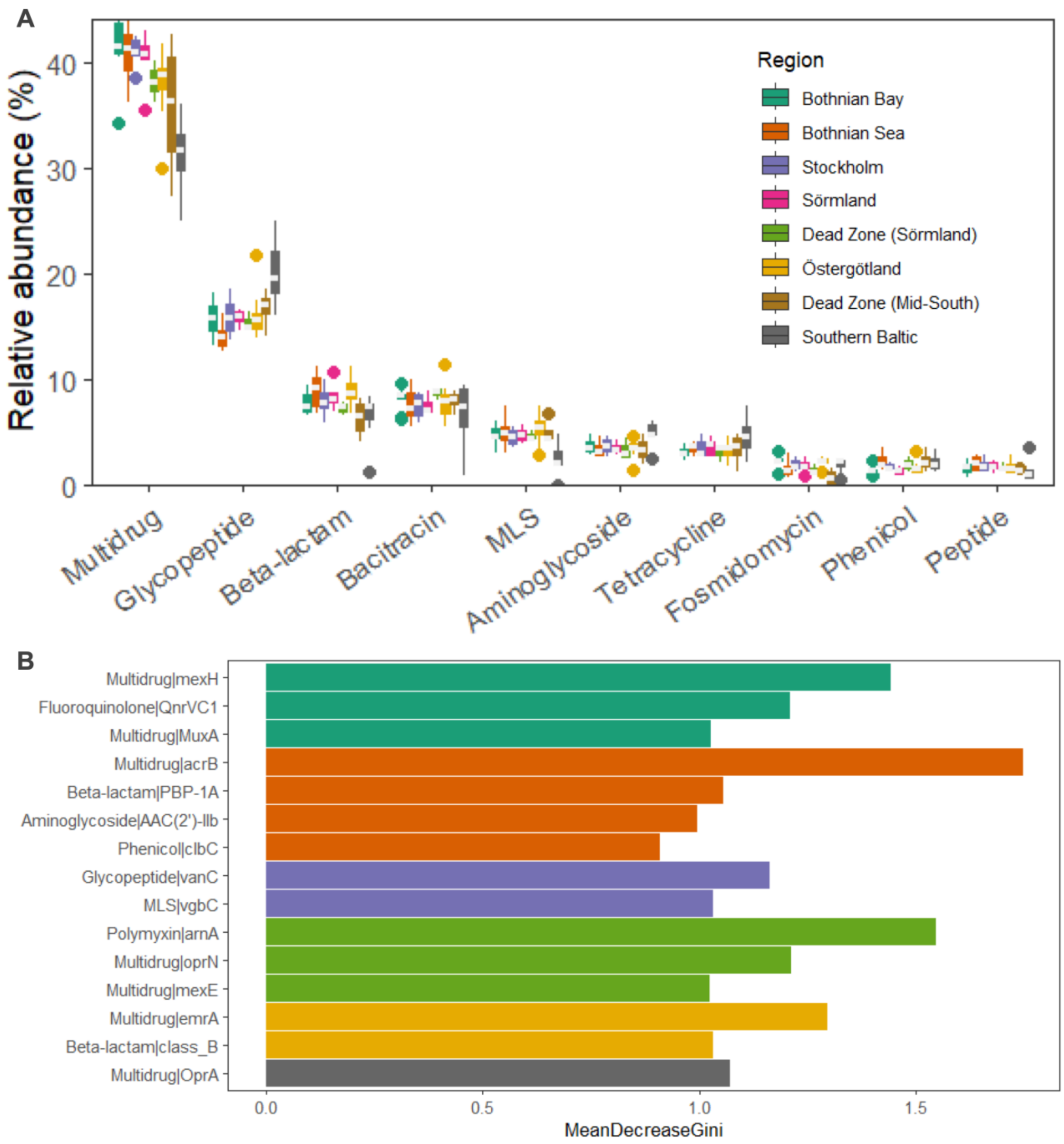


**Supplementary Fig. S5 A** Relative abundance boxplots of the top ARG types across regions. Separate boxplot visualizations of the multidrug and glycopeptide resistance types in Fig. 1D. **B** Differential abundance test by Kruskal-Wallis rank sum and random forest. The variable importance plot shows the top 15 differentially abundant ARG subtypes with high mean decrease Gini scores.

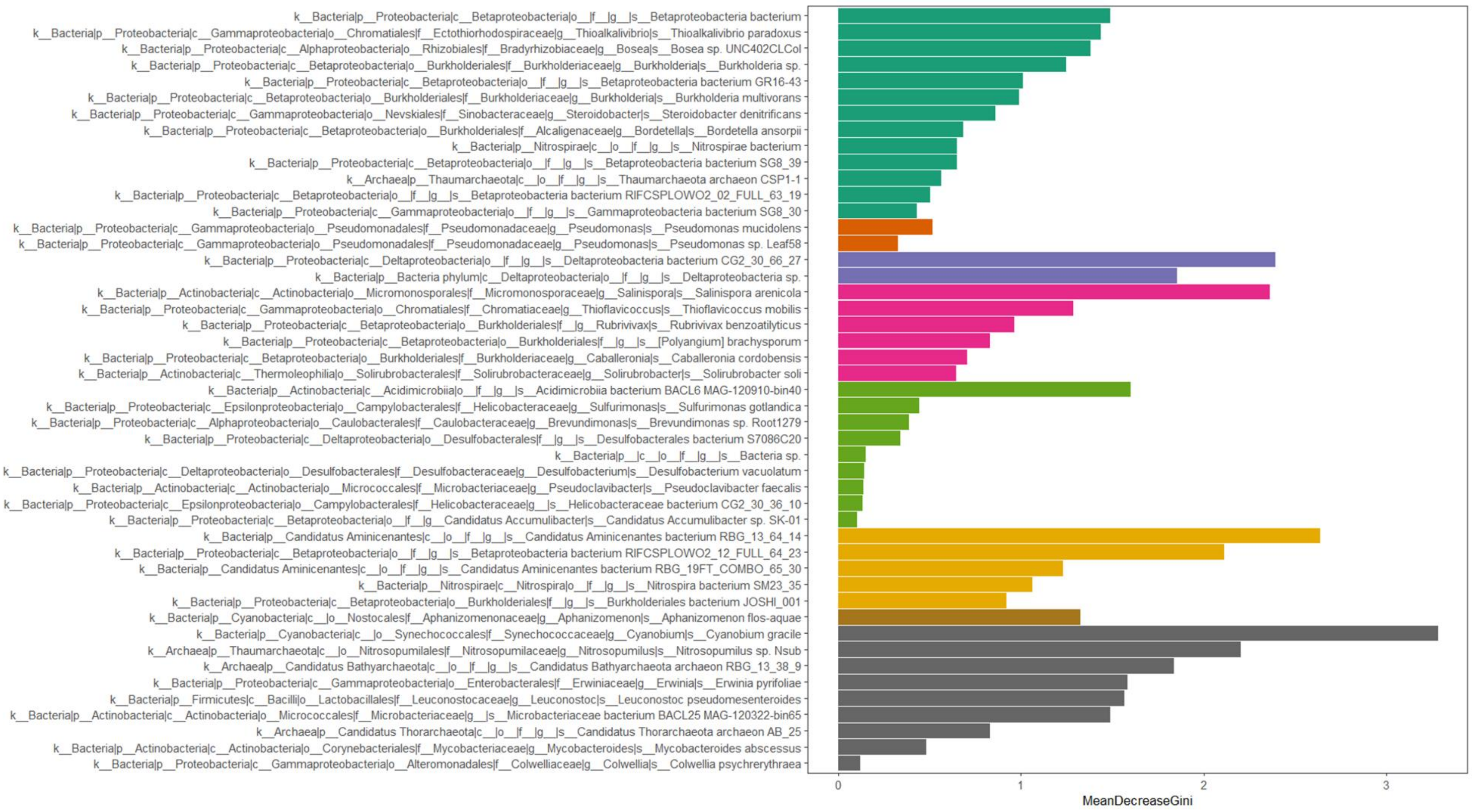


**Supplementary Fig. S6** Differential abundance test by Kruskal-Wallis rank sum and random forest. The variable importance plot shows the 47 differentially abundant species.

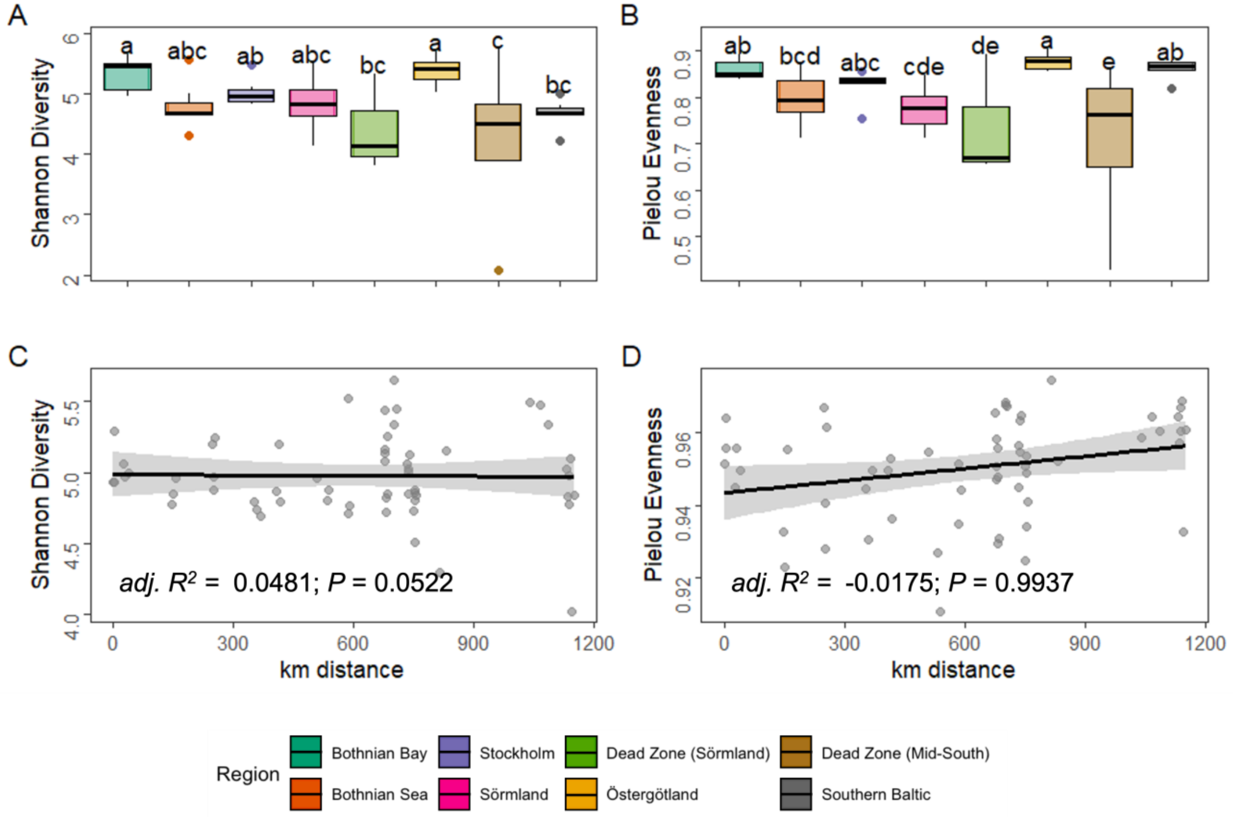


**Supplementary Fig. S7** Alpha diversity of the **A-B** microbial communities and their distribution pattern by **C** Shannon diversity, and **D** Pielou evenness across the Baltic Sea regions in haversine distance (km). Alpha diversity of the **E-G** predicted MRG, **H-I** MGE, and **K-M** VFG. Different letters indicate differences between groups tested by ANOVA or Kruskal-Wallis test (p-value < 0.05).

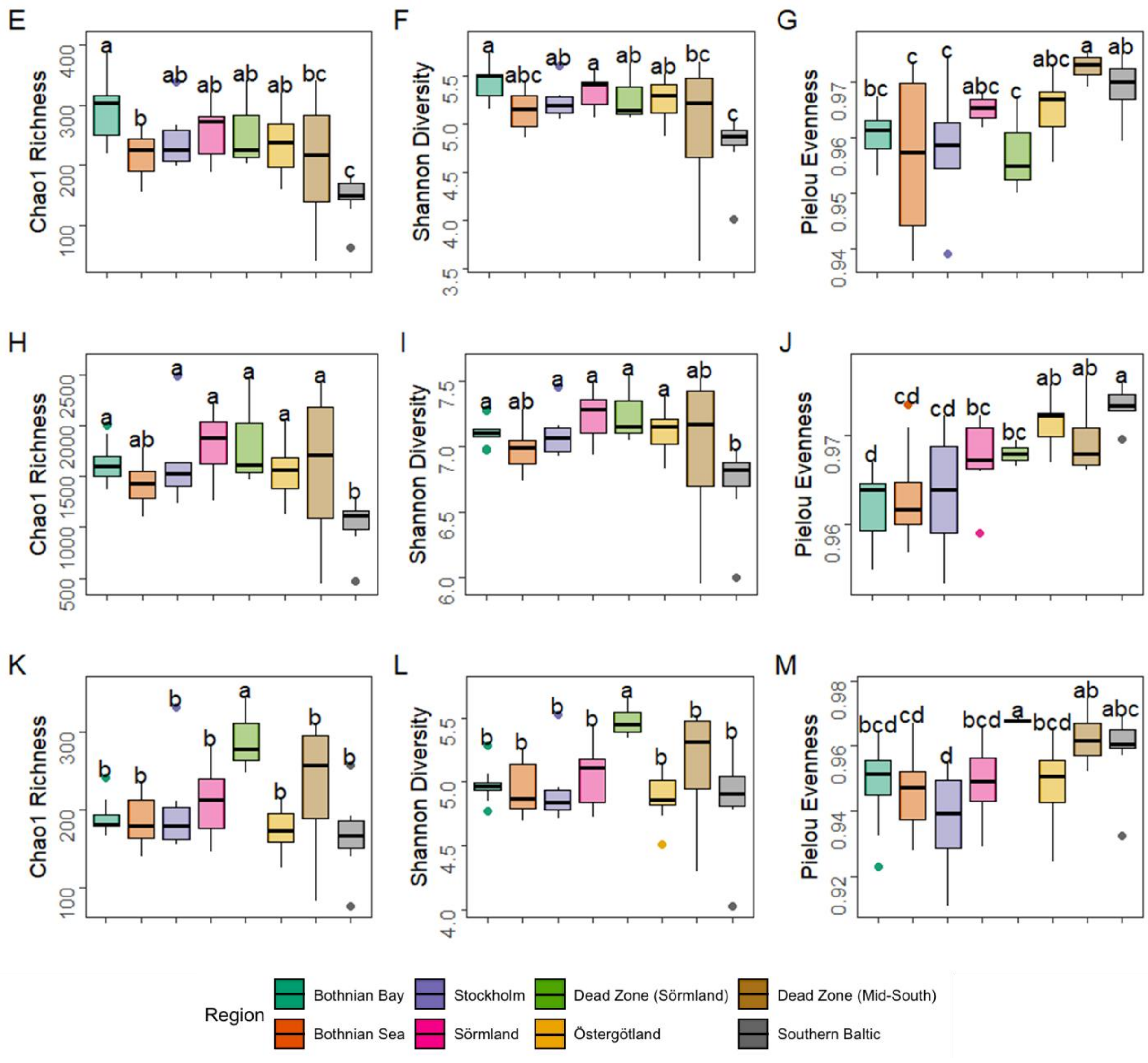


**Supplementary Fig. S7 (cont.)** Alpha diversity of the **A-B** microbial communities and their distribution pattern by **C** Shannon diversity, and **D** Pielou evenness across the Baltic Sea regions in haversine distance (km). Alpha diversity of the **E-G** predicted MRG, **H-I** MGE, and **K-M** VFG. Different letters indicate differences between groups tested by ANOVA or Kruskal-Wallis test (p-value < 0.05).

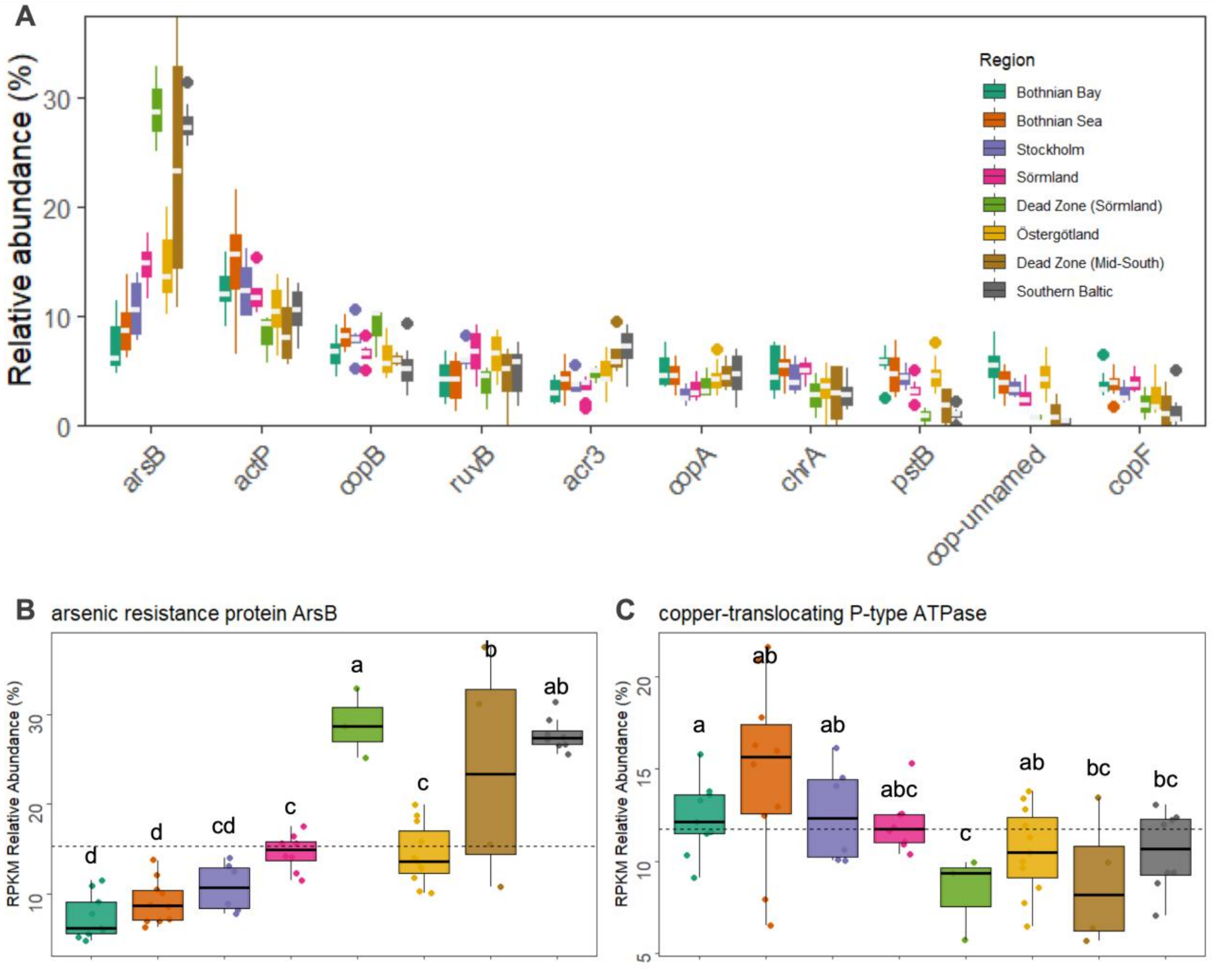


**Supplementary Fig. S8** **A** Relative abundance boxplots of the top MRG types across regions. Separate boxplot visualizations of the **B** arsenic resistance protein *ArsB* and **C** copper-translocating P-type ATPase. Different letters indicate differences between groups tested by ANOVA (P-value < 0.05) .

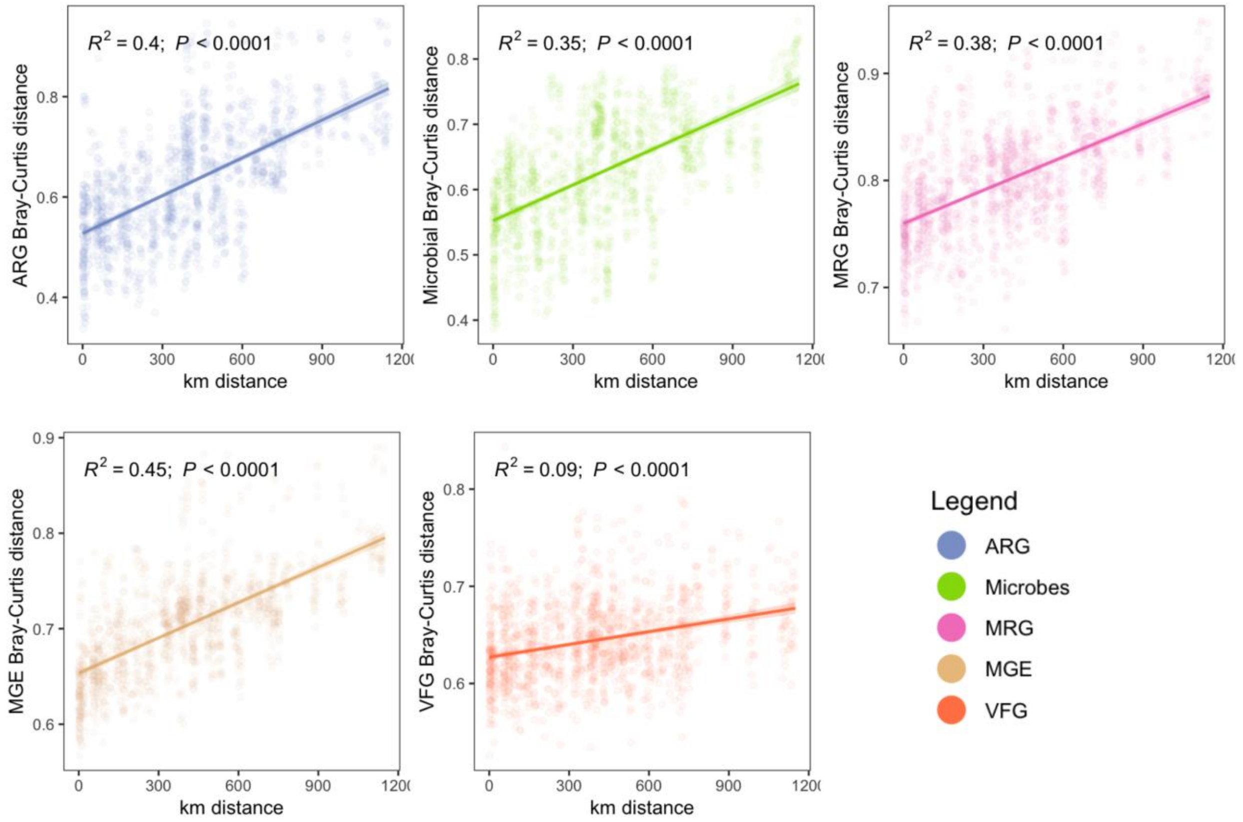


**Supplementary Fig. S9** Distance decay relationship of the Bray-Curtis dissimilarities of the ARG, microbial community, MRG, MGE, and VFG profiles with haversine distance (in km) in the Baltic Sea without the dead zones samples.

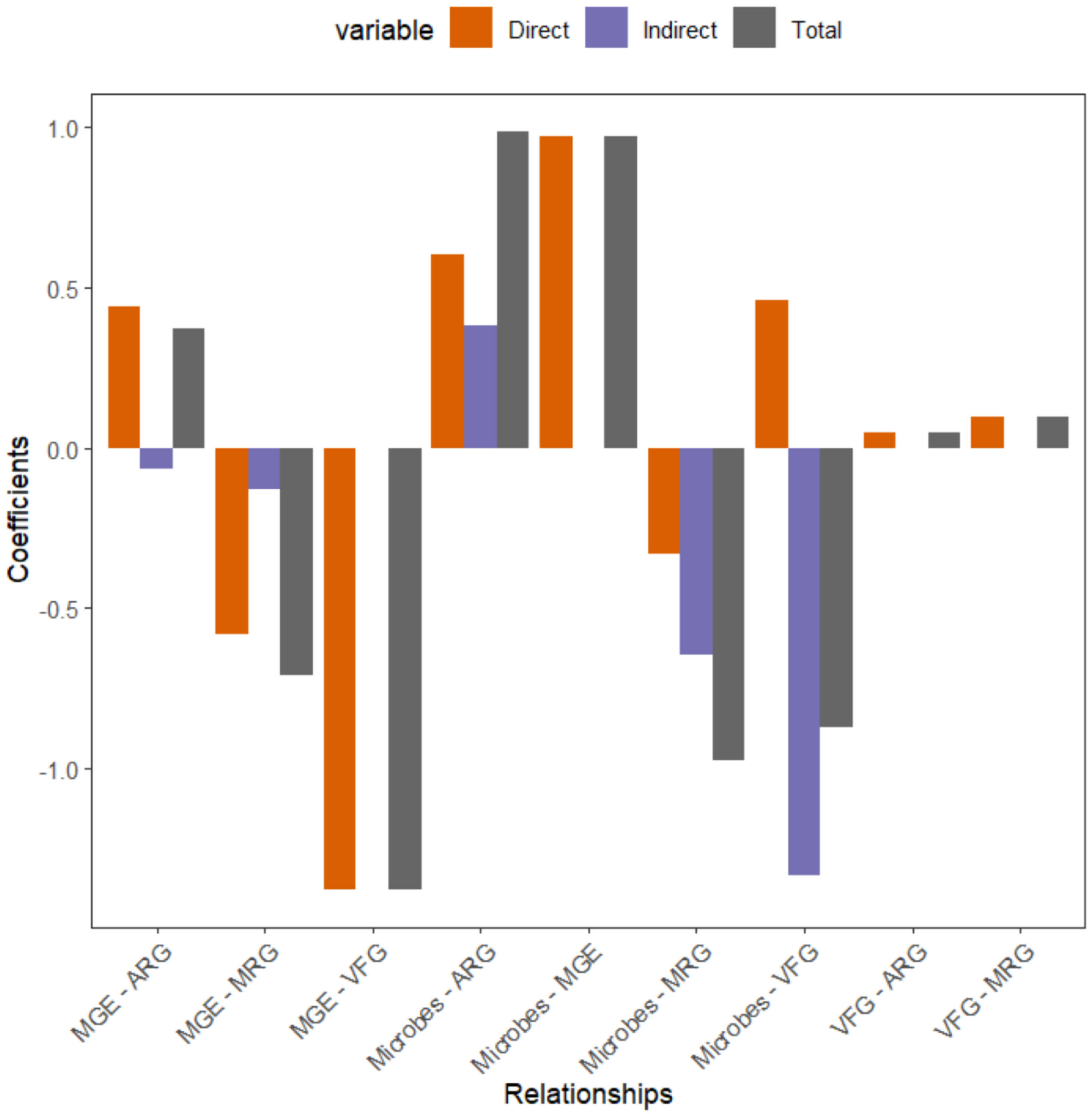


**Supplementary Fig. S10** Effects coefficients of the PLS-PM model of the microbial community, ARG, MRG, MGE, and VFG profiles as latent variables.
